# Hippocampal memory reactivation during sleep is correlated with specific cortical states of the Retrosplenial and Prefrontal Cortices

**DOI:** 10.1101/2023.06.11.544473

**Authors:** PA Feliciano-Ramos, MJ Galazo, H. Penagos, MA Wilson

## Abstract

Episodic memories are thought to be stabilized through the coordination of cortico-hippocampal activity during sleep. However, the timing and mechanism of this coordination remain unknown. To investigate this, we studied the relationship between hippocampal reactivation and slow-wave sleep UP and Down states of the retrosplenial cortex (RTC) and prefrontal cortex (PFC). We found that hippocampal reactivation are strongly correlated with specific cortical states. Reactivation occurred during sustained cortical UP states or during the transition from UP to Down state. Interestingly, sustained UP states from the PFC were more coordinated with memory reactivation in the hippocampus, whereas hippocampal reactivation was biased to occur during the cortical UP to Down state transition of the RTC. Reactivation usually occurred within 150-200 ms of a cortical UP-state onset, indicating that a build-up of excitation during cortical UP state activity influences the probability of memory reactivation in CA1. Conversely, CA1 reactivation occurred 30-50 ms before the onset of a cortical Down state, suggesting that memory reactivation affects Down state initiation in RTC and PFC, but the effect in RTC was more robust. Our findings provide evidence that supports and highlights the complexity of bidirectional communication between cortical regions and the hippocampus during sleep.

## Introduction

The theory of system consolidation proposes a two-stage process for memory formation and stabilization, where information is first encoded during active behavior and then consolidated during sleep (G. Buzsáki 1989; McClelland, McNaughton, and O’Reilly 1995; Kitamura et al. 2017; Klinzing, Niethard, and Born 2019; Stickgold 2005). This consolidation process relies on cortico-hippocampal communication (McClelland, McNaughton, and O’Reilly 1995; MacPherson and Della Sala 2019; Tse et al. 2007; Sawangjit et al. 2018; Kitamura et al. 2017), which has prompted the characterization of timing relationships between cortical and hippocampal sleep oscillations as vehicles of memory transfer and stabilization (Ji and Wilson 2007; Peyrache et al. 2009; Siapas and Wilson 1998; Varela and Wilson 2020; Rothschild, Eban, and Frank 2017; Sirota et al. 2003; Wang and Ikemoto 2016; Qin et al. 1997). While earlier efforts focused on the coupling between hippocampal sharp wave ripples (SWR, 100 - 250 Hz) and cortical spindles (7-15 Hz) or delta (1–4 Hz), the emphasis in recent work has shifted to investigate the relationship between SWRs and cortical slow oscillations (SO, 0.1 - 1Hz) motivated by three important observations. First, SO groups faster sleep brain rhythms including spindles and delta into the complex wave patterns characteristic of slow wave sleep (SWS, Contreras and Steriade, 1995). Second, SO reflects the alternation between Up states, or periods of sustained cortical spiking, and Down states, epochs of little or no cortical activity. Third, hippocampal spiking activity exhibits a similar Up/Down oscillatory pattern during sleep that is temporally linked to the cortical SO despite lacking intrinsic conductances to generate this Up/Down alternation pattern independently. Together, these observations raise the possibility that cortical Up states may serve as computational units that process information transfer from the hippocampus to support memory consolidation and other sleep-dependent cognitive processes (Ji and Wilson 2007; Penagos, Varela, and Wilson 2017). It follows that understanding SWR-SO timing relationships can shed light on these sleep operations.

Importantly, the memory consolidation process is thought to involve bidirectional communications between the cortex and hippocampus during sleep. While the influence of cortical SOs on hippocampal SWRs has been well documented, several questions remain unanswered. For instance, do hippocampal SWRs only respond to SO activity or do SWRs also modulate the features of SO? In addition, because of the strong correlation between SWRs and the reactivation of neuronal ensembles corresponding to waking experiences, a phenomenon termed hippocampal replay (Wilson and McNaughton 1994; Kudrimoti, Barnes, and McNaughton 1999; Foster 2017; Lee and Wilson 2002; Pavlides and Winson 1989; Skaggs and McNaughton 1996), SWRs have been used as a proxy for memory reactivation during consolidation. Although this is supported by observations that selective elimination of SWRs during sleep leads to severe impairments in memory performance (Girardeau et al. 2009; Ego-Stengel and Wilson 2010; Aleman-Zapata, Morris, and Genzel 2022; Aleman-Zapata, van der Meij, and Genzel 2022), the relationship between SO and SWR-associated hippocampal replay is yet to be characterized. Despite studies simultaneously monitoring large-scale cortical dynamics and SWRs (Karimi Abadchi et al. 2020; X. Liu et al. 2021; Nitzan et al. 2022; Logothetis et al. 2012) showing diverse coordination between SWRs, most of these studies lack a behavioral component that would enable exploring the relationship between hippocampal replay and SO in multiple cortical regions.

To address these gaps in knowledge, we performed simultaneous extracellular recordings in the dorsal CA1 layer of the hippocampus and two cortical regions: the retrosplenial (RTC) and prefrontal cortex (PFC). Our choice of these cortical areas stems from their close functional relationship with the hippocampus. PFC is involved in decision-making and rule learning while RTC has been hypothesized to serve as a hub of information to many cortical regions since it is one of the few cortical areas that receive projections from the dorsal hippocampus (Yamawaki et al. 2019; Thomas van Groen and Michael Wyss 1990; T. van Groen and Wyss 1992; Wyass, Michael Wyass, and Van Groen 1992; Ferreira-Fernandes et al. 2019; Opalka and Wang 2020). These close associations have motivated studies in which PFC and hippocampal interactions have been well investigated(Peyrache et al. 2009; Tang et al. 2017; Jadhav et al. 2016; Carr, Jadhav, and Frank 2011; Eichenbaum 2017; Wierzynski et al. 2009) and recent work has also started to characterize how RTC and hippocampus communicate around SWR events (Pedrosa et al., n.d.; Opalka et al. 2020; Pedrosa et al. 2022; Karimi Abadchi et al. 2020; Nitzan et al. 2020; Hale 2015; Alexander et al. 2018; Chambers, Berge, and Vervaeke 2022). We reason that investigating how hippocampal reactivation interacts with RTC and PFC during sleep could help us better understand how hippocampal-related information is broadcast within distinct cortical networks whose interaction is essential to consolidation. To this end, we took advantage of multisite tetrodes recordings and analyzed the local field potentials (LFP), multiunit activity (MUA), and neuronal ensembles to focus on studying the underlying coordination of CA1 memory reactivation with slow wave sleep (SWS)-related UP- and Down-states in the RTC and PFC.

Consistent with previous reports, we found SWRs to be highly correlated with cortical Up states of RTC and PFC (Siapas and Wilson 1998; Sirota et al. 2003; Varela and Wilson 2020; Karalis and Sirota 2022; Opalka et al. 2020). In addition, we also found that SWRs are more likely to occur before a Down state in both areas, with a stronger effect in RTC. Notably, only a minor portion of SWRs occurred during or immediately following a Down-state. Mirroring these findings, CA1 replays mostly occurred during sustained Up states or at the transition from UP to Down states, with stronger coordination observed between RTC Down states and hippocampal replays. Interestingly, both replays and SWRs were less likely to occur during a Down state or in the Down to UP state transition, consistent with the notion that cortical activity drives hippocampal reactivation and not the other way around. Our results support the hypothesis that bidirectional communication between the hippocampus and cortical networks occurs during SWS-dependent memory consolidation. We propose that cortical activity during Up states is a major driver of hippocampal memory reactivation, thus promoting the transfer of information from the hippocampus. Complementing the bidirectional relationship, CA1 reactivation is essential for transitioning from an UP state to a Down state. This, in addition to the role of replay in enabling the exchange between hippocampal and cortical systems, may suggest that the promotion of cortical Down states could be an important component in the consolidation processes.

## Results

### A. Classification of cortico-hippocampal interactions

Cortico-hippocampal information exchange could be optimal when they have synchronized activity (Fries 2005; Penagos, Varela, and Wilson 2017). Consistent with this idea, recent research has shown that hippocampal sharp wave ripples (SWRs) can be coordinated with many brain regions (Nitzan et al. 2022; Karalis and Sirota 2022). Nevertheless, there is a lack of characterization regarding the interactions between cortical UP/Down states and SWRs. Some have reported that SWRs may precede or follow delta-related cortical Down states (Sirota et al. 2003; Battaglia, Sutherland, and McNaughton 2004) or that UP states may contain none, single, or multiple SWRs (Ji and Wilson, 2007); however, it remains unclear which types of interactions are more common within multiple cortical networks. To address this gap in knowledge, we employed a classification approach based on the temporal relationship of cortical UP and Down states with hippocampal SWRs in the CA1 region. We investigated the interactions between hippocampal CA1 and cortical activity in the RTC and PFC during slow-wave sleep (SWS) (Fig. 1, Supplementary Fig. 1A, and Fig. 2). We defined cortical UP or Down states as regions of high or low population cortical activity lasting for 100-1000 ms or 50-500 ms, respectively (Saleem et al. 2010; Karalis and Sirota 2022). Detection of Up and Down states was based on the distributions of the envelope of the high-frequency component (MUAe, multi-unit activity envelope) of the local-field potentials (LFP) or the multi-unit activity (MUA) from detected spikes (Supplementary Fig. 1B A-C). Similar to Ji and Wilson (2007), we observed a bimodal distribution in the MUA and MUAe of the RTC and PFC, representing the UP and Down cortical periods (Supplementary Fig. 1B A-C). When recording cortical activity with a limited number of units (less than 5), MUAe was employed because of its sensitivity in detecting cortical states despite the low levels of spike detection.

**Figure 1:**
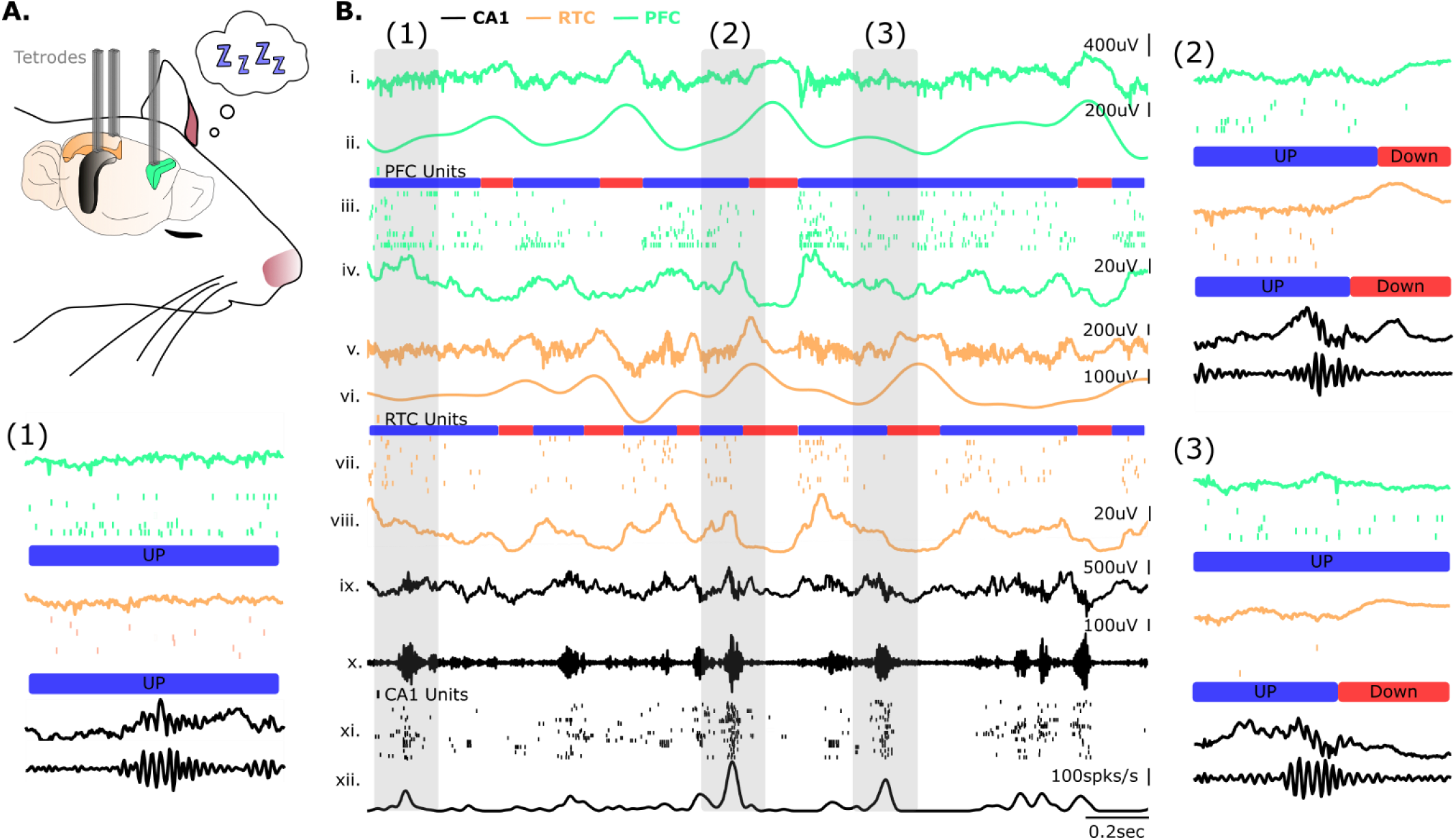
Dynamic coordination of cortico-hippocampal activity during slow-wave sleep (SWS). **A**. Simultaneous extracellular recordings were made in the hippocampal CA1(black), retrosplenial cortex (RTC), and prefrontal cortex (PFC) during sleep of freely behaving animals. Tetrodes (gray) were placed using one of the three configurations: CA1-PFC, CA1-RTC, or CA1-RTC-PFC. The black area in this illustration corresponds to the hippocampal CA1, the orange region to the RTC, and the green region to the prefrontal cortex. **B**. An example of a simultaneous triple extracellular recording between the CA1, RTC, and PFC. i-iv, PFC raw local field potential (LFP), filtered delta (1-4Hz), units, and multiunit envelope (high pass filter > 500Hz). v-viii, RTC raw LFP, filtered delta(1-4Hz), units, and multiunit envelop (high pass filter > 500Hz). x-xii, CA1 raw LFP, units, and multiunit activity. (1), An example of a CA1 sharp-wave ripple (SWR) occurring during cortical activity. (2), An example of a CA1 SWR occurring close to a Down-state region in the PFC and RTC. (3), An example of a CA1 SWR occurring close to an RTC Down-state while PFC was in a UP cortical state.

**Figure 2:**
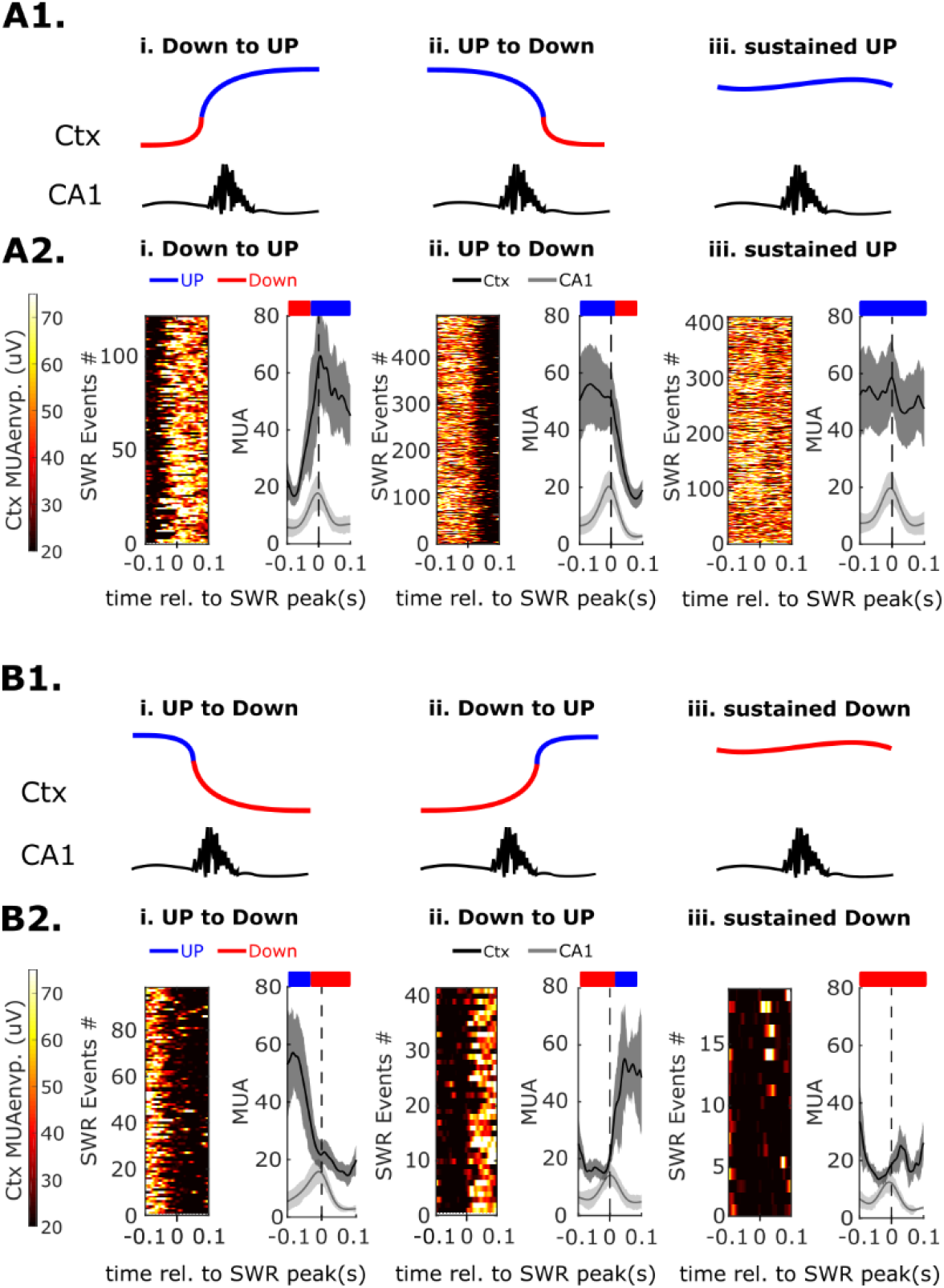
Classification of cortico-hippocampal activity. **A**. Classification of SWRs that occurred during a cortical UP-state. A1, Cartoon illustration of the different SWRs classifications in the UP-state. A2, MUA from the cortex and hippocampus for SWRs classified into different classifications. Left panel, MUAe from the cortex. Right panel, average MUA from cortex and CA1. Red, shows a region of a Cortical Down-state. Blue, a region of cortical UP-state. i-SWRs occurring in the Down to UP cortical state transition. ii-SWRs occurring in the UP to Down cortical state transition. iii-SWRs occurring during a sustained UP-state which is characterized by the lack of detection of a Down-state within the 200ms time-window of analysis. **B**. Classification of SWRs that occurred during a cortical Down-state. B1, Cartoon illustration of the different SWRs classifications in the Down-state. B2, MUA from the cortex and hippocampus for SWRs classified into different classifications. Left panel, MUAe from the cortex. Right panel, average MUA from cortex and CA1. i-SWRs occurring in the UP to Down cortical state transition. ii-SWRs occurring in the Down to Up cortical state transition. iii-SWRs occurring during a sustained Down-state which is characterized by the lack of detection of a UP-state within the 200ms time-window of analysis.

Supplementary Figure 1B C1-C2 shows two representations of a detected Down state from RTC or PFC. In these examples, it can be noticed the reduction in MUA and MUAe during low cortical activity (Down state) followed by a sudden increase in neuronal activity (UP state). We tested the reliability of our detection method for cortical Down states by analyzing their occurrence as a function of the animal’s position and velocity. In agreement with previous studies, Supplementary Figure 1B D1-D2 shows that Down states occur in periods of immobility (low-speed Supplementary Fig1B, D2) and sleep (POST-task Supplementary Fig. 1B, D1-D2), confirming that our method could reliably detect cortical Down states.

Once the Down and UP states were identified, the next step in the analysis involved the categorization of SWRs according to whether they occurred during a cortical UP or Down state. To execute this classification, a 200 ms time window of cortical MUAe or MUA was inspected. This window extended 100 ms before and after the peak amplitude of an SWR event identified in the filtered CA1 LFP (125-300Hz) (Fig. 2). We sorted SWRs into those that happened during UP (Fig. 2A) or Down-state (Fig. 2B) based on a threshold of cortical MUAe or MUA. These two sets were then sub-categorized as follows: Up state to down state transition, SWRs occurring right before the onset of a cortical Down state(Fig. 2Aii or Fig. 2Bi); Down state to Up state transition, SWRs occurring right after the end of a cortical Down state (Fig. 2Ai or Fig. 2Bii); “sustained” UP state, SWRs occurring in the middle of an Up state (Fig. 2Aiii) or sustained Down state, SWRs occurring in the middle of a Down state (Fig. 2Biii) period.

### B. Hippocampal SWRs are more likely to occur during specific RTC and PFC states

SWRs tend to coincide during cortical UP states (Ji and Wilson 2007; Varela and Wilson 2020; Karalis and Sirota 2022; Siapas and Wilson 1998; Kajikawa et al. 2022; Rothschild, Eban, and Frank 2017; Karimi Abadchi et al. 2020; X. Liu et al. 2021), and we wanted to test whether our classification method could replicate these findings. Our results showed that SWRs in the dorsal CA1 were indeed more likely to occur during cortical UP states of the retrosplenial cortex (RTC) and prefrontal cortex (PFC). Moreover, our methodology allowed us to estimate the overall occurrence of SWRs during each cortical state, which was previously unknown. Specifically, the fraction of SWRs occurring during a RTC cortical UP state was 0.828 (IQR: 0.791-0.871), and 0.1724 (IQR: 0.1289-0.2091) for Down states. In the PFC, the fraction was 0.919 (IQR: 0.894-0.934) to 0.08 (IQR: 0.066-0.106) for UP and Down states, respectively (Fig. 3A). Interestingly, when compared to the PFC, the fraction of SWRs occurring during UP or Down states of the RTC was either smaller or larger (UP, *p*=0 95%CI [-0.131, -0.042]; Down, *p*=0 95%CI [0.041, 0.131] robust ANOVA post hoc bootstrap-t). To determine whether this coordination occurred by chance, we randomly shuffled the SWRs in time and ran the classification algorithm with the same parameters as in the experimental conditions. Our results showed that the proportion of SWRs occurring during UP and Down states was statistically different from the shuffle in both regions (Supplementary Fig. 3 A1 and B1). This shuffle analysis aligns with prior research indicating that cortical UP states predominate during SWS. Furthermore, the statistically significant distinction between the observed proportion of SWRs and the shuffled condition implies that the temporal coordination between SWRs and cortical UP and Down states in the RTC and PFC is not random.

**Figure 3:**
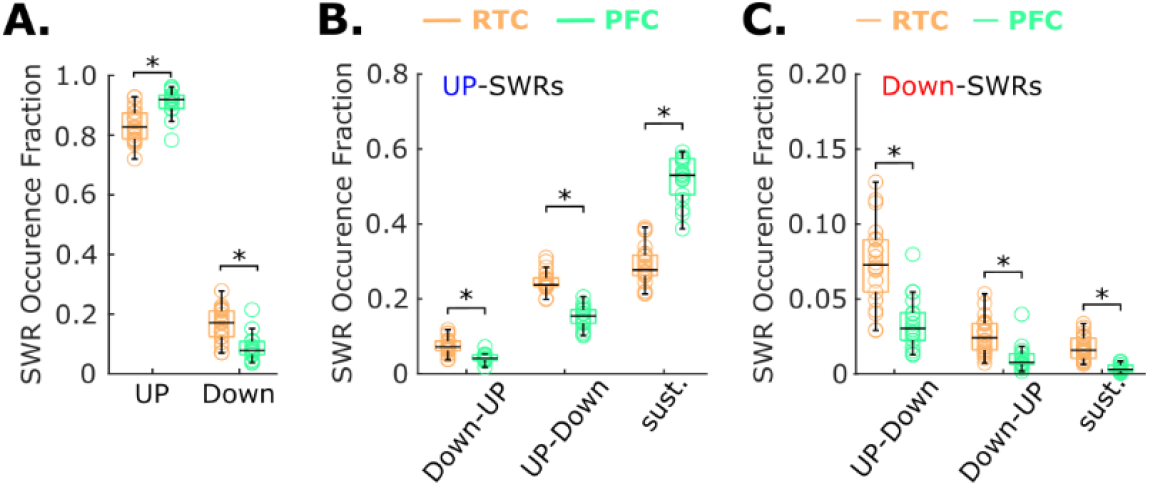
Classification and coordination of hippocampal CA1 SWR activity with the RTC and PFC cortical UP and Down states. **A**, Fraction of SWRs occurring during a cortical UP or Down-state of the RTC or PFC. **B**, Fraction of SWRs occurring during different classifications of the UP-state. States transition from Down to Up, Up to Down, or sustained (sust.) UP-states. **C**, Fraction of SWRs occurring during different classifications of the Down-state. States transition from Up to Down, Down to Up, or sustained Down-states. RTC n=22 8 mice; PFC n=18 5 mice; *=p<0.01 robust ANOVA with post hoc bootstrap-t.

This classification method offers another advantage by expanding its ability to characterize a broader range of temporal coordination, an aspect that has not been explored in previous research. Building on this, we conducted further investigations into the interactions between SWRs and cortical states, as depicted in Fig. 1 and 2. Our findings revealed that most SWRs occurred during specific cortical UP state configurations, as illustrated in Fig. 3B-C. Among the various types of coordination, the most frequent was found during the sustained UP-states of RTC or PFC, as shown in Fig. 3B. Our analysis involving shuffling demonstrated that this particular configuration was also the most likely to occur (Supplementary Fig. 3A2-B2). However, the occurrence rate in the experimental condition was significantly higher than that in the shuffles for both RTC and PFC, indicating that this type of coordination is not random. Additionally, we observed that the fraction of SWRs during sustained UP-states was greater in the PFC than in the RTC (Fig. 3B, sust.-UP PFC = 0.667 (IQR: 0.615-0.721) vs sust.-UP RTC = 0.349 (IQR: 0.331-0.398), p= 0.0, 95%CI [-0.370, -0.248], robust ANOVA post hoc bootstrap-t). Showing that SWRs in CA1 are more likely to be coordinated with sustained cortical UP state activity in the PFC than in the RTC.

We observed that the second most frequent classification in our analysis was for SWRs occurring during a cortical UP state, followed by a Down state (Fig. 2 Aii and Fig. 3B). These SWRs were classified as UP to Down events (Fig. 2Aii, Fig. 3B). The fraction of SWRs during this transition was 0.299 (IQR: 0.294-0.321) for RTC and 0.195 (IQR: 0.170-0.215) for PFC. This coordination was found to be more frequent in the RTC compared to the PFC (Fig. 3B UP-Down, *p*= 0.0, 95%CI [0.085, 0.143], robust ANOVA post hoc bootstrap-t). Furthermore, when examining SWRs occurring during the Down state (Fig. 3C) and in the UP to Down transition configuration (Fig. 2Bi), we discovered that the median fraction for the RTC was higher than that for the PFC. Specifically, RTC had a median fraction of 0.072 (IQR: 0.056-0.088), whereas PFC had a median fraction of 0.030 (IQR: 0.023-0.040) (Fig. 3C, *p*= 0.0, 95%CI [0.023, 0.059], robust ANOVA post hoc bootstrap-t). Interestingly, this classification is similar to the UP to Down configuration observed during the UP state, with the only difference being a slight temporal shift in the occurrence of SWRs. Taken together, these results highlight that SWRs in CA1 have stronger temporal coordination with cortical Down states of the RTC compared to the PFC. This emphasizes a notable difference in the coordination patterns between RTC and PFC.

Although the results presented above describe the most frequent classifications, they do not provide information on the degree of coordination. To address this, we calculated the ratio between experimental and shuffle classification fractions (Supplementary Fig. 3C1-C2), which served as a metric to assess the degree of dissimilarity between experimental classification outcomes and a random process. As expected, the configurations in the UP-state, such as the Up to Down (RTC 1.950 (IQR: 1.860-2.297); PFC 1.405 (IQR: 1.316-1.564)) and sustained UP (RTC 1.632 (IQR: 1.527-1.747); PFC 1.433 (IQR: 1.360-1.486)), were the only configurations that showed ratio values larger than one (Supplementary Fig. 3A C1). This indicates that SWRs are strongly coupled to periods of sustained cortical activity and to UP to Down state transitions. Remarkably, among all configurations, the UP to Down configuration of RTC displayed the highest ratio values (Supplementary Fig. 3A C1). Although this type of coordination was not the most frequent, the analysis indicates that the interaction between SWRs and RTC during the UP to Down transition represents the most temporally coordinated pattern.

Lastly, our classification method revealed that SWRs occurred less frequently during the Down to Up transition of the UP or Down state and during sustained Down states (Fig. 3B-C). What the three configurations have in common is that a Down state precedes SWRs, indicating that a lack of activity from the cortex dramatically reduces the probability of SWR occurrence. To further investigate the low fraction numbers observed in these configurations, we examined whether our analysis failed to detect them and tested the reliability of the shuffle analysis. If SWR occurrence and cortical states were independent random processes, we would expect equal probability for SWRs occurring before or after a cortical Down state. Consistent with this notion, we found that the fraction of SWRs in the shuffled data occurring during the UP to Down or Down to Up transitions in both RTC and PFC were similar, regardless of whether SWRs were associated with Up (Fig. 2Ai-ii; *p=* 0.9 95%CI [-0.016, 0.015] for RTC, and *p*= 0.94 95%CI [-0.018, 0.018] for PFC, robust ANOVA post hoc bootstrap-t) or Down (Fig. 2Bi-ii; *p*= 0.868 95%CI [-0.021, 0.019] for RTC, and *p*= 0.734 95%CI [-0.014, 0.011] for PFC, robust ANOVA post hoc bootstrap-t) states. Furthermore, all these configurations had a fraction value smaller than their respective shuffles (Supplementary Fig. 3 A2-A3, B2-B3, C1-C2). This analysis allows us to draw several conclusions. First, the lower fraction of Down to Up and all Down configurations, compared to their respective shuffles, indicates that the low occurrence of these configurations is not due to limitations in the detection method. Second, the shuffle analysis successfully captured the independent nature of the shuffled data, indicating that the coordination of SWRs and cortical states in each configuration is well coordinated and far from random interaction.

### C. The temporal coordination between SWRs and Down states is stronger in RTC than in PFC

In this section, we focused on identifying the reasons behind the higher likelihood of certain cortico-hippocampal configurations, particularly those where SWRs occurred during the transition from UP to Down, which is the most coordinated configuration between hippocampal CA1 and RTC (Supplementary Fi3A C1). To determine this, we examined the onset of Down states in RTC and PFC and CA1 SWRs delta phase distributions (Fig. 4B). Prior studies have established a correlation between delta oscillations and cortical Down states during SWS. Therefore, we first analyzed whether the onset of detected Down states was coupled with the delta phase. Our results showed that the delta phase distribution for the onset of Down states of RTC and PFC was phase coupled (Figure 4B Rayleigh’s test *p*<0.01). The average delta phase distribution for the onset of Down states of the RTC was 145° and for PFC was 120°. Next, we examined the phase distribution of hippocampal SWRs to the delta wave of RTC and PFC. The delta phase distribution for SWR with respect to delta in RTC was 150°, and for PFC, it was 30°. As previously reported, SWRs in CA1 are temporarily phase coupled to delta waves from RTC and PFC. However, our analysis uncovered an interesting observation regarding the absolute differences between the SWR-delta phase and the onset of delta phase distribution of Down states. Notably, these differences were smaller in the case of RTC compared to PFC (Fig. 4B3, *p*=0 95%CI [-77.646 -37.848], robust t test bootstrap-t). This indicates that SWRs and the onset of delta-related the UP to Down transition in RTC, as depicted in Figure 3B.

**Figure 4:**
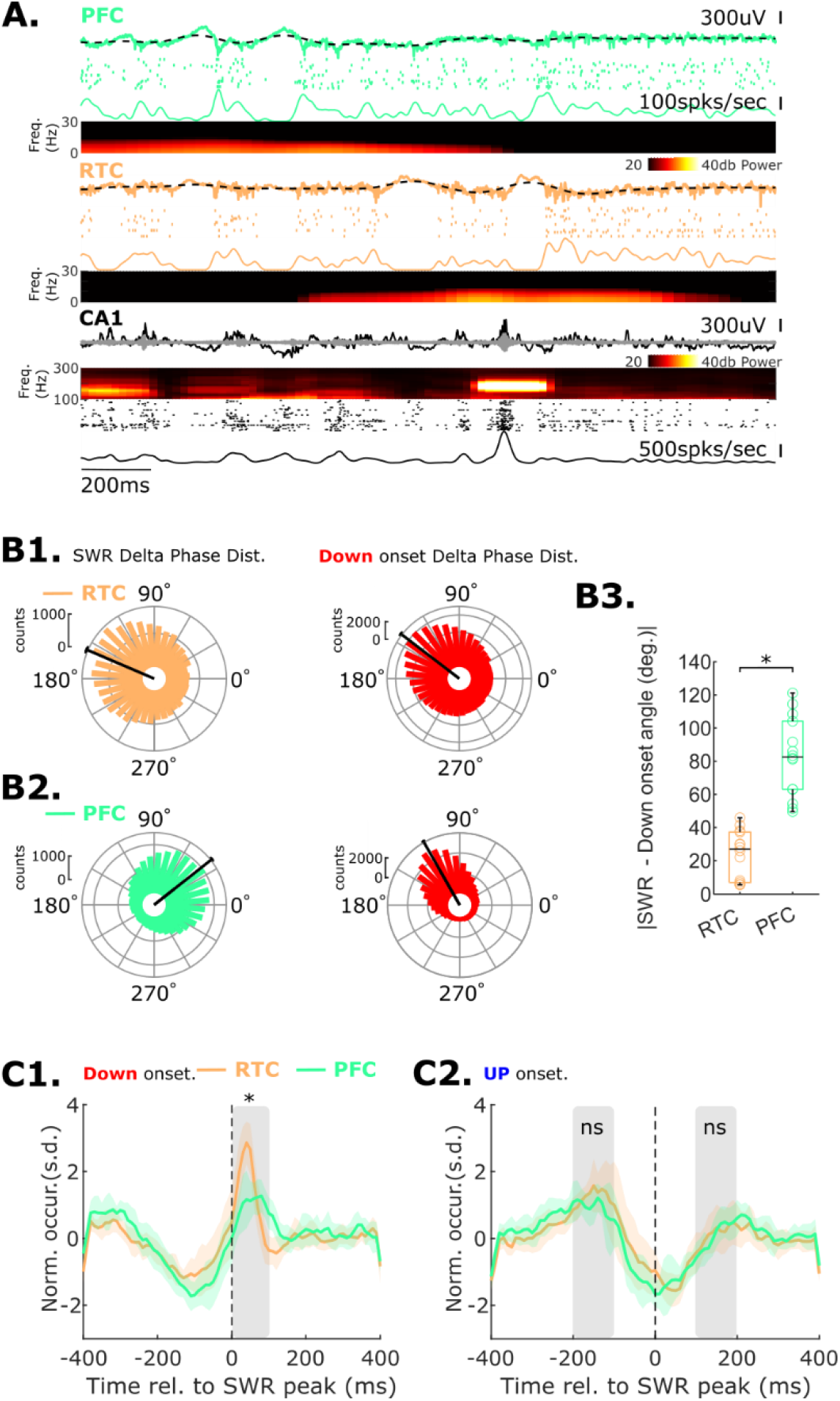
SWRs and Down states delta phase distributions and UP/Down states onset peri-event time histograms. **A**, An example of a simultaneous triple extracellular recording between the CA1, RTC, and PFC. PFC raw local field potential (LFP), filtered delta (1-4Hz black), units, and MUA firing rate, spectrogram. RTC raw LFP, filtered delta (1-4Hz black), units, and MUA firing rate, and spectrogram. CA1 raw LFP, filtered ripples (125-300Hz grey), spectrogram, units, and MUA. **B**, SWRs and Down states delta phase distributions. B1, RTC SWRs (orange) and Down states (red) delta phase distributions. Mice 4, n=14. B2, SWRs and Down states delta phase distributions. B1, PFC SWRs (green) and Down states (red) delta phase distributions. Mice 4, n=14. **C**, Peri-events time histograms for RTC and PFC Down (C1) and UP (C2) states onsets during SWRs. *= *p*< 0.01, ns= *p*>0.05. Only simultaneous triple recordings included in the analysis.

Next, we investigated the possibility of hippocampal SWRs increasing the probability of cortical Down states of RTC and PFC. To do so, we performed a peri-event time histogram analysis on cortical UP and Down states that occurred during SWRs (Fig. 4C). We examined a time window of 800 ms (400 ms before and after the peak of SWRs) and looked at the onset of cortical Down states in RTC and PFC (Fig. 4C1). We found that the probability of a Down state onset increased shortly after the appearance of an SWR, with the lowest point occurring 200-150 ms before the SWR and then suddenly increasing after the SWR. This aligns with previous findings in PFC and RTC. However, we observed a greater peak probability of Down state onset in RTC than in PFC, indicating stronger coordination between hippocampal SWRs and RTC Down states onset. The z-score median peak amplitude of a Down state onset in RTC was twice as large as that in PFC (RTC, 3.087 (IQR: 2.934-3.525); PFC, 1.413 (IQR: 1.178-1.939); Fig. 4C1 and Supplementary Fig. 4A1, *p*=0 95%CI [1.269 2.033], robust t test bootstrap-t). To calculate the variance and timing of the peri-event histogram between 0 and 100 ms (gray region in Fig4 C1) after an SWR peak, we fitted a Gaussian. Consistent with our previous finding that the delta phase distributions for RTC and SWRs were more closely aligned than the phase distributions between SWRs and PFC, we found that the onset of a Down state peaked at a median of 28.89 ms (IQR 26.35-32.40ms) for RTC and 44.12ms (IQR 41.55-48.37ms) for PFC (Fig 4C1 and Supplementary Fig. 4A3; *p*=0 95%CI [-19.040 -11.382], robust t test bootstrap-t). Additionally, we noted a decrease in the variance of the onset of Down state in RTC compared to PFC, with a variance of 38.44 (IQR 35.50-44.91) for RTC and 54.55 (IQR 51.73-85.22) for PFC (Supplementary Fig. 4A2; *p*=0 95%CI [-52.615 -12.478], robust t-test bootstrap-t). A decrease in variance is also an indication that the initiation of RTC Down states is more Down states in RTC tend to occur in closer phases compared with PFC. This finding provides a partial explanation for the higher occurrence of SWRs during tightly linked to SWRs in CA1 than Down states in PFC. To ensure that these results were not influenced by variations in the quality of the Gaussian fit, we conducted a comparison of the root-mean-square deviation (RMSE) for each fit. The median RMSE values were similar for both groups, with 0.490 (IQR: 0.376-0.631) for RTC and 0.508 (IQR: 0.424-0.667) for PFC (Supplementary Fig. 4A4, p=0.68, 95% CI [-0.197, 0.122] robust t test bootstrap t).

Additionally, we investigated whether the stronger coordination between SWRs and Down states from RTC was specific to Down states. To examine this, we analyzed the peri-events histogram for the onset of cortical UP states in RTC and PFC. Our observations revealed that SWRs typically occurred around 150-200 ms before or after an SWR event (Fig. 4C2). Notably, the likelihood of an UP state onset before an SWR was slightly higher compared to the peak amplitude after an SWR (Fig. 4C2). For RTC, the median zscore peak amplitude for an UP state onset before and after a SWR was 2.289 (IQR: 1.790-2.598) and 1.2405 (IQR: 0.519-1.371), while for PFC was 1.786 (IQR: 1.626-1.931) and 0.907 (IQR: 0.795-1.300), respectively (RTC *p*=0 95%CI [0.519 1.939]; PFC *p*=0.005 95%CI [0.413 1.155] robust ANOVA post hoc bootstrap-t). However, there was no difference in the occurrence of UP state onset before or after an SWR event when comparing RTC and PFC (Fig. 4C2 gray areas, Supplementary Fig. 4B1-B2, RTC vs PFC Before *p*=0.114, 95%CI [-0.092 0.969], RTC vs PFC After *p*=0.972 95%CI [-0.545 0.533] robust ANOVA post hoc bootstrap-t). Based on the higher probability of an RTC or PFC UP state occurring before an SWR, our results support the notion that cortical UP state activity can influence SWR occurrence. Furthermore, our findings indicate that SWRs do not appear to affect the initiation of cortical UP states in RTC and PFC, but rather they may primarily impact Down state initiation. Thus, a higher probability of an UP state preceding an SWR serves as an indicator of SWR occurrence, while an SWR serves as a reliable indicator of a cortical Down state occurrence, with a more pronounced effect observed in the RTC.

### D. CA1 replays occur during specific cortical states of RTC and PFC

Next, we examined the relationship between memory reactivation (replays) in the hippocampus and its interaction with cortical states of the RTC and PFC. The model for the system consolidation suggests that during sleep, the hippocampus replays neural activity patterns from wakefulness, which is essential for the memory consolidation. Our experimental findings indicate that the most favorable forms of interaction between SWRs in the CA1 region and the cortical states of the RTC and PFC occur either during sustained UP states in the cortex or during the transition from UP to Down. However, it is unclear whether hippocampal replays follow the same timing with respect to cortical states.

To answer this question, we recorded place cells in the hippocampal CA1 while mice were running in a linear maze of approximately 200 cm (Fig. 5A1). We then used Bayesian decoding to detect spatial trajectories from the spiking activity of hippocampal place cells during sleep. To validate our approach, we created an encoding model using 80% of the data while the animal was running in the linear maze and subsequently predicted the animal’s position on the remaining 20% of run data, based on CA1 spiking activity. Figure 5 A2 and A3 show the confusion matrix and decoding error cumulative distribution of a single experimental session demonstrating our ability to estimate the mice’s position on the maze (median error 4.81 cm). Next, we applied the model to the spiking activity during sleep to detect replays. First, we selected periods of elevated spiking activity (Davidson, Kloosterman, and Wilson 2009; Wu and Foster 2014; Chen and Wilson 2017; S. Liu, Grosmark, and Chen 2018) as candidate replay events. Using the maximum likelihood estimator (MLE) at each time point (Davidson, Kloosterman, and Wilson 2009; Wu and Foster 2014; Chen and Wilson 2017; S. Liu, Grosmark, and Chen 2018), we assessed the spatiotemporal structure of each candidate event by applying linear and weighted distance correlations between MLEs and time. The resulting correlation values were assessed for statistical significance by comparing them to distributions of spatio-temporal correlations obtained from randomly shuffled spiking data (Davidson, Kloosterman, and Wilson 2009; Wu and Foster 2014; Chen and Wilson 2017; S. Liu, Grosmark, and Chen 2018; Tingley and Peyrache 2020).

**Figure 5:**
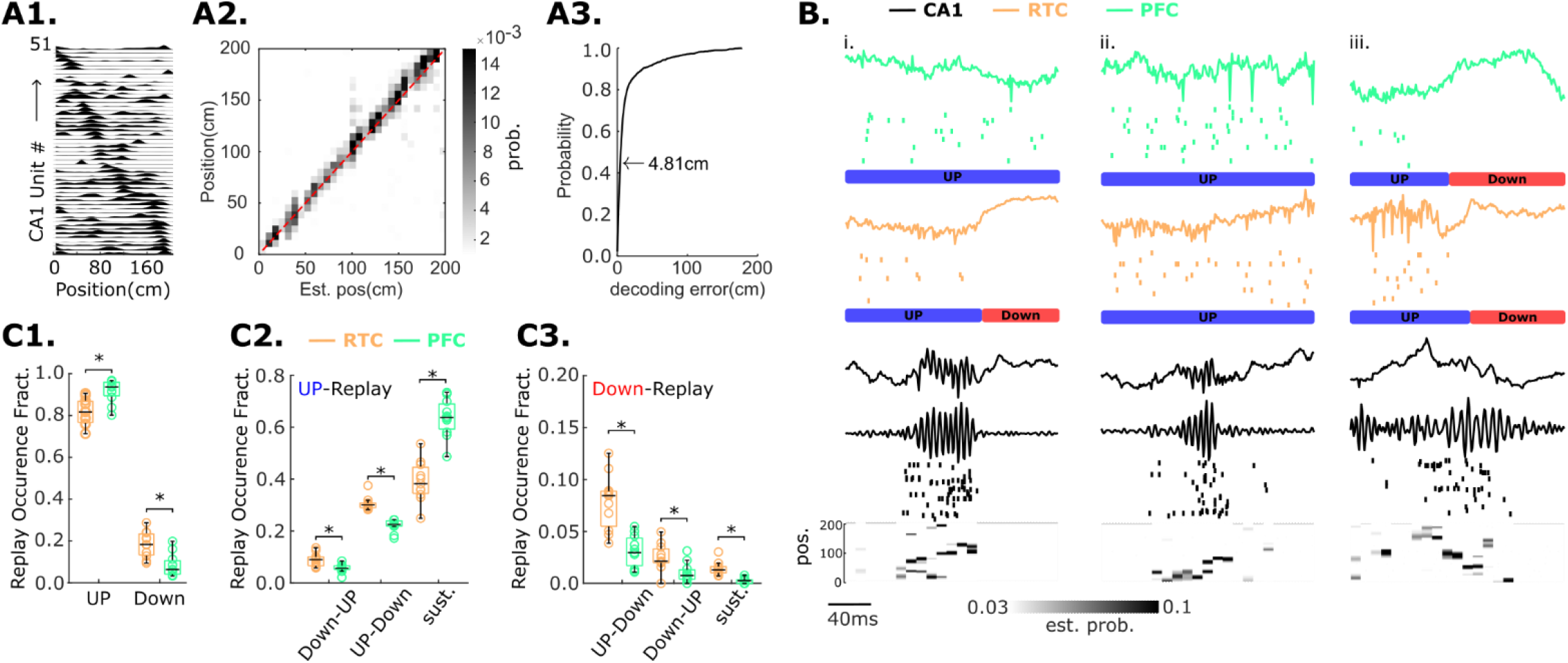
Coordination of hippocampal Replays and cortical states of the RCT and PFC. **A**, Bayesian decoding using place tuning units from hippocampal CA1, A1, Firing rate of CA1 Units in a linear maze. Units are ordered based on their place field. A2, Confusion matrix showing the distribution of the actual position of a mouse and the estimated position from the decoding model. A3, Empirical cumulative distribution function (CDF) of decoding errors. The arrow represents the median error in estimating the position for a single animal experiment. **B**, Replay examples and their coordination to cortical states of the RTC and PFC, In green the LFP and units of PFC, In orange the LFP and units of RTC, In black the LFP, filtered LFP ripples, units, and replay position reconstruction of CA1; An example of a replay that was coordinated with an UP to Down transition in RTC, but in PFC, it remained in a sustained UP state. ii-An example of a replay coordinated to sustained UP states of the RTC and PFC. iii-An example of a replay coordinated to a UP to Down transition in RTC and PFC. **C**, Classification and coordination of hippocampal Replays with RTC and PFC cortical UP and Down states. C1. Fraction of replays occurring during a cortical UP or Down-state of the RTC or PFC. C2, Fraction of replays occurring during different classifications of the UP-state. States transition from Down to Up, Up to Down, or sustained (sust.) UP-states. C3, Fraction of replays occurring during different classifications of the Down-state. States transition from Up to Down, Down to Up, or sustained Down-states. RTC n=11, 3 mice; PFC n=11, 2 mice; *=p<0.01 robust ANOVA with post hoc bootstrap-t.

In Figure 5B, we show a sample of three replayed sequences detected in CA1 using our replay detection method. To study replay interaction with cortical states of the RTC and PFC, we employed the same classification method to investigate the fraction of replays that occurred during the UP and Down states of the RTC and PFC. Consistent with our earlier findings showing a higher probability of SWRs occurring during cortical UP states, we observed that replays were more likely to occur during cortical UP states than during Down states of the RTC and PFC (Fig5 C1, RTC UP vs Down, *p*=0 95%CI [0.523 0.732]; PFC UP vs Down *p*=0 95%CI [0.788 0.927] robust ANOVA post hoc bootstrap-t). Hippocampal replays occurring during an UP state of RTC had a median occurrence fraction of 0.817 (IQR 0.773 -0.863) and for PFC, it was 0.936 (IQR 0.901-0.955). Conversely, for RTC, the median fraction of replays occurring during Down states was 0.183 (IQR 0.137-0.227) and for PFC was 0.064 (IQR 0.045-0.099). These results provide evidence that hippocampal reactivation is strongly coordinated with UP states of multiple cortical regions such as RTC and PFC.

Next, we investigated other types of coordination between memory reactivation from hippocampal CA1 and the cortical states of RTC and PFC. As previously found, hippocampal sequences during sleep are less likely to occur during the transition from the Down to Up state of RTC or PFC. The median fraction of replays occurring during this transition was 0.086 (IQR 0.066-0.097) for RTC and 0.053 (IQR 0.049-0.062) for PFC, making this configuration the least likely of the UP state configurations (Fig. 5C2). Furthermore, replays in CA1 during Down states have the lowest probability among all configurations. For example, replays occurring during sustained Down states have the lowest occurrence probability with a median fraction of 0.013 (IQR 0.010-0.016) for the RTC and 0.003 (IQR 0.000-0.003) for the PFC. The second-lowest occurring probability was for replays occurring during the Down to Up transition of a Down state, with a median fraction of 0.021 (IQR 0.020-0.031) for the RTC and 0.008 (IQR 0.004-0.012) for the PFC (see Fig. 5C3). Interestingly, these two configurations have the second largest window of cortical inactivity of all classifications. In contrast, the Down state configuration where replays occur in the transition from Up to Down, which is the configuration where cortical activity precedes the occurrence of a replay, was the most frequent configuration in this group (Fig. 5C3). The median occurrence fraction for these replays was 0.085 (IQR 0.059-0.088) for the RTC and 0.029 (IQR 0.021-0.042) for the PFC (see Fig. 5C3). This indicates that periods of cortical inactivity greatly reduce the probability of hippocampal reactivation.

Finally, and similar to our previous results, we found that the most likely coordination between hippocampal replays and the cortex occurs during the UP state configuration, such as the sustained UP states and in the transition from UP to Down of the RTC and PFC (Fig5 C2). In particular, we found that the median fraction of replays occurring during sustained UP states was stronger in PFC (0.6335, IQR 0.6006-0.6755) compared to RTC (0.3784, IQR 0.3463-0.4311) (see Fig. 5C2, *p*=0 95%CI [-0.368 - 0.137] robust ANOVA post hoc bootstrap-t), indicating that memory reactivation from hippocampal CA1 is more likely to interact with PFC and RTC during periods of prolonged cortical activity, but this coordination is stronger between replays and PFC. The second most probable configuration for replays was during the transition from UP to Down of RTC and PFC, with a median fraction of occurrence of 0.297 (IQR 0.288-0.303) for RTC and 0.222 (IQR 0.217-0.232) for PFC (Fig. 5C2). Interestingly, replays showed stronger coordination with the UP to Down transition of RTC compared to PFC (Fig. 5C2, *p*=0 95%CI [0.050 0.094] robust ANOVA post hoc bootstrap-t). It is worth noting that in both configurations, cortical activity preceded hippocampal replays, indicating a loop-like coordination between the cortex and hippocampus. Our data provide evidence that prolonged or sustained periods of cortical activity increase the likelihood of memory reactivation, whereas extended periods of cortical inactivity significantly reduce the probability of hippocampal SWRs and replay occurrences. Additionally, we observed that memory reactivation in the hippocampus triggers the transition from the UP to Down state in both RTC and PFC. These findings support the notion of a bidirectional interaction between the cortex and hippocampus, where hippocampal reactivation serves dual purposes: transmitting sequences to the cortex and promoting the initiation of the Down state, the latter likely serving as a mechanism to minimize memory interference.

## Discussion

Here we report coordinated activity between hippocampal sharp wave-ripples (SWRs) in CA1 and the retrosplenial cortex (RTC) and prefrontal cortex (PFC) during slow-wave sleep (SWS). We employ a novel classification method to investigate distinct patterns of communication between SWRs and the RTC, and PFC. Our findings demonstrate that SWRs and their interaction with the cortical activity are not random but consistent with the idea of two-way communication between these regions. Hippocampal SWRs and replays are more likely to occur during specific cortical states such as sustained cortical activity (UP-states) or during periods in the state transition from UP to Down. One consistent observation across these configurations is that cortical activity precedes SWRs and memory reactivation in the hippocampus. The probability of a replay or SWR occurrence is higher after 150-200 ms following the initiation of a cortical UP state. In contrast, SWRs or replays occurring during prolonged Down states or in the state transition from Down to Up are much less frequent, indicating that periods of inactivity greatly reduce the probability of SWRs and memory reactivation. We hypothesize that cortical input into the hippocampus is a major driver of hippocampal reactivation. Additionally, we observed that SWRs and replays often occur during the cortical state transition of UP to Down of the RTC and PFC. The onset of a Down state of the RTC and PFC occurs within 30-50 ms after the peak of SWRs. In contrast to the occurrence of the onset of a cortical UP state, the initiation of a Down state before an SWR shows an oscillatory behavior that is disrupted after the appearance of an SWR. This suggests that SWRs and replays can provide feedback input into cortical regions such as the RTC and PFC by locally increasing the probability of a Downstate. Importantly, hippocampal feedback seems to have a different impact depending on the cortical region because the coordination of Down state onsets and the fraction of SWRs and replays occurring during the state transition of UP to Down is greater between the hippocampus and RTC.

### A. Differences in RTC and PFC cortical state interactions between SWRs and replays

Our findings revealed several interactions between hippocampal reactivation and RTC and PFC. Specifically, during sustained UP-states, we observed that this was the most frequent type of interaction between SWR/replays and PFC and RTC. Notably, the frequency of these interaction was higher for the PFC compared to the RTC, challenging the prevailing notion that slow oscillations (SOs) during slow-wave sleep (SWS) are uniformly synchronized across all cortical regions. Our findings highlight the complexity and diversity in the coordination of SWRs and replays with the cortical state activity of the PFC and RTC. We propose that this interplay is not solely reliant on synchronized activity but involves dynamic flexibility, allowing for selective coordination. This intricate interplay could enables the hippocampus to transmit information to the cortex through various mechanisms.

The second most frequent type of interaction in our classification method was during the state transition from UP to Down of the RTC and PFC. SWRs preceding a cortical Down state (UP to Down) have been reported in several cortical regions such as the sensory cortex(Sirota et al. 2003), PFC (Karalis and Sirota 2022; Peyrache et al. 2009) and in RTC (Opalka et al. 2020). In agreement with those studies, we also see strong coordination between SWRs and cortical Down states of RTC and PFC; however, we are the first to report that this coordination is more robust between the hippocampus and RTC. Moreover, our shuffle analysis demonstrated that SWRs occurring in the UP to Down configuration of RTC was the most coordinated interaction despite not being the most frequent. We reason that memory reactivation in the hippocampus will exert a higher cortical influence in areas of the RTC than in PFC with respect to promoting downstate generation. Global Up/Down states are thought to be generated by the thalamocortical and cortico-cortical networks, likely playing a key role in the coordination of various cortical regions (Connelly et al. 2015; Vaasjo et al. 2022; Narikiyo et al. 2020). Here, our study reveals that the hippocampus has the capacity to locally coordinate Down states in both the PFC and RTC, with a more significant impact observed in the RTC. We reason that locally coordinating activity in cortical regions through hippocampus inputs could be an important mechanism information processing.

### B. A build-up of excitation from the cortex and circuit dynamics are responsible for the coordination of SWRs and replays during cortical UP states

Our research findings show that memory reactivation in the hippocampus is strongly coordinated with cortical UP states of the RTC and PFC, which may be attributed to anatomical and circuit dynamic interactions. We observed a delay of 150-200 ms before SWRs and replays reached their peak occurrence probability, which explains the higher fraction of SWRs/replays occurring during sustained UP-state configurations in our classification method. The delay in SWRs and replays occurrence is in line with the idea that a build-up of cortical excitation may be responsible for the increased probability of memory reactivation in the hippocampus. This may also explain why SWRs and replays are less likely to be observed immediately during the onset of cortical UP states (Down to UP transition) or during Down state configurations. However, the anatomical complexity of brain regions must also be taken into consideration, as information needs to travel and interact through different circuits before promoting SWRs and memory reactivation in the CA1 and therefore contributing to the observed delay in our results.

Interestingly, recent evidence from simultaneous intracellular and extracellular recordings in hippocampal CA1, CA3, dentate gyrus (DG), and cortex has revealed that a complex and lengthy process could ultimately trigger the occurrence of SWRs in CA1 (Kajikawa et al. 2022). Even though the mechanism for SWR initiation is not known, experimental evidence suggests that hippocampal regions from CA2 and CA3 are important in promoting SWRs and replays in CA1 (Yamamoto and Tonegawa 2017; Oliva et al. 2016; Ecker et al. 2022; de la Prida 2020; Davoudi and Foster 2019). However, recent research conducted by Kajikawa et al. (2022) highlights the significance of cortical input in coordinating hippocampal activity. Their study found that cortical UP states play a crucial role in modulating the membrane potential of DG, CA3 and CA1 neurons. Interestingly, the study observed that in CA1, the cell membrane potential is depolarized about 100 ms prior to the initiation of SWRs, where the depolarization period was coordinated with the cortical UP state. This observation is particularly important because the duration of this depolarization coincides with our finding that the peak occurrence probability of a cortical UP state occurs approximately 150 ms before/after the occurrence of an SWR. Based on these results, prolonged and coordinated interactions between cortical and hippocampal pathways are critical for initiating SWRs in CA1. We reason that the cortex mostly mediates the buildup of excitation in the hippocampus, which in conjunction with circuit dynamics, plays an essential role in memory reactivation within the hippocampus.

### C. The probability of SWRs and replays occurrence is low during cortical Down states

The hypothesis of circuit dynamics and build-up excitation is useful in explaining the strong coordination between SWRs and replays to cortical UP states. However, this does not explain why a minority of SWRs and replays occur during cortical Down states. Replays during Down states are believed to be instances where the hippocampus leads the transfer of information (Nitzan et al. 2020; Karimi Abadchi et al. 2020). Our study found that the probability of coordinated activity where hippocampal SWRs precede cortical UP states is low. Nevertheless, we recognize that this coordination may still be crucial for memory consolidation. Experimental and computational evidence indicates that hippocampal-hippocampal connections have the circuit capacity to generate SWRs and replays without the need for external input (György Buzsáki 2015). This could indicate that SWRs observed during Down states may be generated by internal hippocampal circuits such as from CA3 and CA2 in which without cortical input they may represent a group of replays where their content is purely influenced by internal hippocampal computations.

Alternatively, it is plausible that these SWRs and replay events that occurred during Down states of the RTC or PFC are triggered by other cortical regions, and as a result, they may not represent a “leading” scenario for the hippocampus. For example, the medial entorhinal cortex (MEC) is known to undergo spontaneous persistent activity (SPA), a phenomenon that allows the MEC to skip UP/Down state cycles from neocortical regions (Hahn et al. 2012; Choudhary et al. 2022; Egorov et al. 2002; Yoshida, Fransén, and Hasselmo 2008). If the MEC skips a neocortical Down state(e.g. from RTC or PFC) and stays in UP state instead, it could increase the likelihood of SWRs and replays in CA1 that occurred during Down states of the RTC or PFC. The average occurrence rate of SPA in MEC neurons is around 5-20%, which is similar to the overall occurrence rate of replays and SWRs that we reported during Down states. Although we are uncertain about the extent to which SPA in the MEC could influence memory reactivation in the hippocampus, research has shown that optogenetic inhibition of MEC inputs to CA1 can decrease the probability of SWRs and replay initiation (Yamamoto and Tonegawa 2017). Therefore, we believe that the low numbers of replays that occur during cortical Down states of RTC or PFC are influenced by other cortical regions such as the MEC. This hypothesis could further diminish the role of the hippocampus in leading the information exchange to the cortex since, under such circumstances, distinct cortical regions could be the ones dictating the timing of hippocampal memory reactivation.

### D. Down states of the RTC are strongly coordinated to hippocampal SWRs/replays

The occurrence of Down states in RTC and PFC reaches its peak shortly after an SWR in dorsal CA1, which indicates that the probability of a cortical Down state in RTC and PFC is increased by hippocampal reactivation. Coordination between Down states in the RTC and SWRs/replays was found to be stronger than that observed between Down states in the PFC. This finding is supported by a higher fraction of SWRs and replays occurring in the UP to Down configuration and increased likelihood of a Down-state following an SWR. The reason for the difference in coordination is unclear but may relate to anatomical and circuit dynamics.

The PFC and hippocampal interactions occur through direct and indirect projections from the intermediate and ventral hippocampus to the PFC (Eichenbaum 2017). These projections are both excitatory and GABAergic (Melzer and Monyer 2020). SWRs in the ventral, intermediate, and dorsal CA1 are not synchronous but strong SWRs, which are those with large recruitment of neurons can propagate through the longitudinal axis of the hippocampus (De Filippo and Schmitz 2023; Sosa, Joo, and Frank 2020). Modulation of GABAergic circuits in cortical layers has been involved in the regulation of cortical Down states (Zucca et al. 2017; Narikiyo et al. 2020; Jackson et al. 2018; Fanselow and Connors 2010). We thus hypothesize that certain SWRs in the dorsal CA1 can propagate through ventral and intermediate hippocampal projections that could directly modulate a population of GABAergic cells in the PFC either directly or indirectly via the thalamus, which can then facilitate Down states in the PFC. In contrast, the RTC receives hippocampal projections from the dorsal area, and the mechanism of how they can modulate Down states in the RTC is better understood. Opalka et al. (2018) demonstrated that specific optogenetic stimulation of the axons of the dorsal hippocampus in RTC can cause a biphasic modulation of inhibitory and excitatory neurons from the RTC (Opalka et al. 2020). Either optogenetic stimulation or basal SWRs promoted excitatory cells in the RTC to increase their firing rate, followed by a rapid inhibitory period. The inhibitory cells, on the other hand, were modulated directly but with a delay in their firing rate. This finding suggests that the coordination observed between SWRs and the transition of the cortical state from UP to Down in the RTC is possible due to the robust modulation of RTC inhibitory circuits via hippocampal projections.

In addition to the difference in dorsal vs ventral hippocampal projection to RTC and PFC, a new study has also found more anatomical differences that could account for the difference in the strength of coordinated between Down states of the RTC and PFC with dorsal SWRs. Ferreira-Fernandes et al., (2019) used novel anatomical tracing tools to shed light on the different connectivity patterns between the hippocampus and the medial mesocortex, which includes the cortical regions from RTC to the cingulate cortex (CGC). The study found that the hippocampal projections into the medial mesocortex exhibit a gradient-like connectivity pattern (Ferreira-Fernandes et al. 2019). Specifically, the RTC has denser projections and stronger connectivity between the dorsal and intermediate CA1 compared to the prefrontal areas, which reinforce our observation that anatomical and circuit differences are responsible for the stronger coordination between Down states in RTC and SWRs/replays in the dorsal CA1. However, the consistency of this interaction is not always reliable, and not every time an SWR ends up finishing an UP state. It is important to discuss that it is possible that other brain circuits, such as the thalamus, may be required to be included in this process to increase the probability of this coordination (Tomé, Sadeh, and Clopath 2022). In addition, an alternative hypothesis could be that SWRs and replays, such as those that propagate along the hippocampal longitudinal axis, may have a stronger impact on cortical Down states of the RTC or PFC during certain memory-related processes, particularly during the reactivation of long and extended experiences (Davidson, Kloosterman, and Wilson 2009; Yamamoto and Tonegawa 2017). In support of this idea, studies have shown that long SWRs in the hippocampus, which involve more neurons representing task-relevant information, are critical for learning and memory (Fernández-Ruiz et al. 2019; Ngo, Fell, and Staresina 2020). This could indicate that the hippocampus may enhance the coordination of multiple cortical regions by increasing the likelihood of Down states, particularly during prolonged experiences when many neurons are recruited. This mechanism could play an essential role in consolidation and require further investigation.

### E. Cortical state coordination with hippocampal reactivation during SWS

Cortico-hippocampal interactions are essential for memory consolidation, and we found that the coordination of SWR/replays with cortical states of the RTC and PFC is highly specific and well-coordinated. We were intrigued by the previously overlooked connection between SWR/replays and cortical UP to Down transitions in sleep studies. This strong correlation left us wondering why the hippocampus would influence the transition of the cortex to the Down state after memory reactivation. The current evidence related to cortical Down states and their relationship with SWR/replays is somewhat controversial.

During wakefulness, earlier research has shown that Down states, also known as “micro-sleep” periods, occur more frequently when there is poorer behavioral performance (Vyazovskiy et al. 2011). Recent studies have supported this finding and have utilized advanced closed-loop technology to investigate the relationship between Down states and SWRs during behavior. In particular, these studies have shown that inhibiting the prefrontal cortex after the detection of an SWR can have negative effects on performance (den Bakker et al. 2022; Peyrache et al. 2009). These findings suggest that during wakefulness, the cortical Down state that follows a hippocampal SWR may be inversely related to memory formation. On the contrary, other studies have suggested that SWRs and Down state interactions during sleep could play a vital role in memory stabilization. For instance, Maingret et al. (2016) demonstrated for the first time that increasing the coordination of SWRs with cortical Down states in the prefrontal cortex through electrical stimulation during sleep can improve memory consolidation in a spatial object task (Maingret et al. 2016). Furthermore, Todorova and Zugaro (2019) found that a small population of cortical neurons activated during cortical Down states formed cell assemblies with SWRs-activated neurons during sleep (Todorova and Zugaro 2019). They proposed that the inhibitory action from cortical Down states can be used to reduce non-relevant information during cell assembly formation, as the cells that were correlated with SWRs were selectively modulated during the task. In addition, a recent study by Kim et al. (2023) found evidence that cortical Down states in the motor and PFC, together with hippocampal SWRs, increase their coupling during learning and disengage during adaptation(Kim et al. 2023). This has been proposed as a mechanism that supports the dual-stage hypothesis for memory consolidation, where cross-brain region coupling via SOs with SWRs is essential for the early stage of memory consolidation.

In line with these studies, we reason that the coordination between replays and Down states during the UP to Down transition in cortical regions could play multiple roles during sleep-dependent memory processes such as consolidation. Replays have long been thought to be solely an information transfer mechanism that in coordination with cortical activity is used for establishing neuronal ensembles between cortical-hippocampal modules (Rothschild, Eban, and Frank 2017; Kaefer et al. 2022). In addition to this important mechanism, our data indicate that hippocampal reactivation can promote cortical Down states. The effect that we see relates to the local coordination, where SWRs can strongly modulate certain cortical regions more than others. This could give the hippocampus the flexibility to engage with specific cortical regions during certain memory processes where a specific type of information is needed. We proposed the hypothesis that the excitatory effect resulting from hippocampal memory reactivation is necessary for the formation of memory ensembles between cortical-hippocampal regions. Moreover, when this reactivation is accompanied by the modulation of Down states during SWS, it could enhance the flexibility of the hippocampus to selectively coordinate brain regions or reduce memory interference. Ultimately, these mechanisms may contribute to the successful formation of memory ensembles.

## Material and Methods

### Mice

We used male C57BL/6J mice (n = 9; 25–32 g; 12–16 wk old at the time of surgery for drive implantation) purchased from Jackson Laboratories. After drive implantation, mice were singly housed in a temperature-controlled room with standard mouse cages (29 x 19 x 12.7 cm) that contained bedding (EcoBedding and nestlets) on a 12 h light/dark cycle (lights on/off at 7:00 am/7:00 pm) with ad libitum access to water and food. All experiments were approved by the Committee on Animal Care at the Massachusetts Institute of Technology and conformed to US National Institutes of Health guidelines for the care and use of laboratory animals.

### Surgical Drive implantation and electrophysiology recordings

Multielectrode arrays (microdrives) with up to 16 nichrome wire tetrodes(individual tetrode wires were 12.5 µm in diameter). The microdrive were prepared according to standard procedures in the laboratory, and the base was modified from previous designs (Kloosterman et al., 2009; Nguyen et al., 2009) to target the three regions of interest. Because replay recordings require many units, we placed 10 tetrodes in dorsal CA1, 3 in RTC, and 3 in PFC and used the common average referencing (CAR) plugin from the Open-Ephys software as a reference for all recordings. To ensure a balanced representation of CAR signal and prevent dominance by any single region, channels from all three sites were included while removing any damaged channels before generating CAR. The tetrodes spanned the following coordinates (Paxinos and Watson, 1986): Bregma +2.3 mm to 0.5 mm medial/lateral for mPFC, Bregma -1.7 mm to 0.5 mm medial/lateral for RTC, and Bregma -1.7-2.0 mm to 1-1.75mm medial/lateral for dorsal CA1.

Sterile surgical procedures were performed for chronic microdrive implantation; anesthesia was induced and maintained with 1–2% inhaled isoflurane. Up to 4 bone screws were secured to the skull for support; two Craniotomies were drilled over the target coordinates and the dura mater membrane was removed to allow for tetrode penetration. The implant was secured to the skull with dental cement after surrounding the exposed craniotomies and tetrodes with silicon grease to prevent them from being fixed by the cement. Animals were preemptively injected with analgesics (Buprenorphine, 0.5–1 mg/kg, subcutaneous) and monitored by the experimenter and veterinarian staff for 3 days post-surgery.

The tetrodes were gradually advanced to the target depths over the course of 1 week before recording started. Once the target depth was reached, minimal adjustments (≈ 25-50uM movement) were made the night before to guaranty the recording from different populations of cells and the stability of unit and local field potential (LFP) recordings. Sleep recording sessions typically lasted at least 1.5 h, while behavior and sleep recordings took around 3-4 hours. All sleep recordings were made in a home cage surrounded by walls preventing mice from accessing visual cues.

Spikes and LFPs were acquired at 30 kHz using Intan Technologies RHD-64 recording headstage with a 64-channel amplifier chip and Open-Ephys GUI (https://open-ephys.github.io/gui-docs/index.html). Units were detected and isolated using kilosort2.0 (https://github.com/MouseLand/Kilosort/releases/tag/v2.0) and Phy GUI(https://github.com/cortex-lab/phy). LFPs were downsampled to 1 KHz from up to 30 KHz using the Matlab downsample function. DeepLabCut (https://github.com/DeepLabCut/DeepLabCut) was used to track the mouse position and estimate velocity.

### LFP and event state detection

For each mouse, we selected one wire from one or both cortical regions with clear sleep events (determined by visual inspection) and used 1-2 tetrodes. These selections were made for LFP/unit sleep analyses. To analyze cortical and hippocampal oscillations, we used two LFPs from each region (CA1, RTC, and PFC). We applied a band-pass finite impulse response filter designed with a Blackman window to filter the signals in the delta (1–4 Hz), ripple (125–250 Hz), or high-frequency (100-500 Hz) bands. Zero-phase distortion was ensured during the filtering process.

Sleep periods were identified as regions characterized by high delta power and immobility (velocity < 1.5 cm/sec). The threshold for analyzing sleep periods was determined by examining the bimodal distribution of delta power, which distinguished between awake and sleep states. Subsequently, various analyses such as replays, sharp wave rippleS (SWR) detection, UP/Down states, and shuffle analysis were conducted specifically on the detected sleep periods. Any events detected outside these sleep periods were excluded from the analysis.

SWR events were detected using Hilbert transformed from the filtered ripple band. A 25 ms Gaussian smoothing filter (Matlab smoothdata function) was applied to the filtered signal, and a threshold of 3 standard deviations from the mean was used to identify candidate events. For an event to be considered as an SWR event, it had to meet the criteria of having a duration <25ms and <500ms, with an inter-interval larger than 25 ms.

Cortical UP/Down states were detected using a similar method as described by Ji and Wilson in 2001. The method involved analyzing the bimodal distribution of the multi-unit activity (MUA) or MUA envelope (MUAe) obtained from the high-frequency (100-500Hz) filtered signal of the cortical tetrodes’ local field potentials (LFPs) (Supplementary Fig 1). To derive the MUAe, we applied the Hilbert Transform to the high-frequency band of the LFPs in the RTC and PFC and smoothed the signal using a Gaussian filter with a 25 ms time window. The use of MUAe has been helpful in previous studies when dealing with challenging clustering or spike detection scenarios(Choi et al. 2010; Ahmadi, Constandinou, and Bouganis 2021). The high-frequency component of LFP is a reliable indicator of ionic conductivity from cortical activity and can be useful in detecting UP/Down states, as demonstrated in Supplementary Figure 1B, C1, and C2 (Reimann et al. 2013). Only cortical UP/Down states during sleeping periods (high-delta power) were included in the analysis. For our study, we defined a cortical UP-state as a period of cortical activity represented by MUA or MUAe that lasted longer than 100 ms but less than 1 sec. Conversely, a cortical Down state was defined as a period of low/no cortical activity in the MUA and MUAe that lasted longer than 50 ms but less than 500 ms. It is worth noting that our definition of cortical UP/Down states aligns with the definition reported by Karalis and Sirota in 2022(Karalis and Sirota 2022).

### Cortical State Classification

To classify the cortical state during SWR replays, we start by obtaining the peak amplitude from the ripple band Hilbert transform of the SWR or the peak amplitude of the CA1 MUA for replay classification. This serves as the reference point, and we extend it 100 ms before and after, creating a total time window of 200 ms. For classification, we calculate the median of a 70 ms time window before (−200 to -130ms), during (−35 to 35ms), and after (130 to 200ms) the MUA or MUAe from the RTC and PFC. This results in a matrix size of 3 in the number of ripple/replay events. To determine the cortical state, we compared the calculated median with a threshold. If the median is below the threshold, it is classified as a Down state region. Conversely, if the median is above the threshold, it is classified as an UP-state region. Figure 2 illustrates the classification results for each configuration. In Supplementary Fig. 3B, we present the uncategorized SWRs that occurred between cortical UP states (Supplementary Fig. 3B A1, B1-B2) or Down states (Supplementary Fig. 3B A2, C1-C2). However, due to the duration of the UP and Down states being on average less than 50 ms, these events are faster than delta oscillations. Consequently, it becomes challenging to ascertain whether they represent typical UP or Down states. As a result, we made the decision not to include them in our analysis. The fraction of cortical SWR/replay state classification was calculated by dividing the number of classifications by the total number of SWR/replay events, including the uncategorized ones.

### Delta phase distribution for SWR and Down-states

The phase of the delta wave for RTC and PFC was estimated using the Hilbert transform from the delta band during periods of high delta power (sleeping periods). We obtained the timestamp from the SWR peak amplitude and the onset of a cortical Down state from RTC and PFC to determine the delta angle for SWRs and Down states. The average angle distribution and the Rayleigh test of uniformity were obtained using the Matlab functions CircHist (https://github.com/zifredder/CircHist) and CircStat (https://github.com/circstat/circstat-matlab). Only simultaneous triple recordings (CA1, RTC, and PFC) were included in this analysis.

### UP and Down state onset occurrence during SWRs

A peri-event histogram for the onset of UP and Down states was obtained from a time window of 800 ms, where 0 represented the SWR peak amplitude. The count of events was then z-scored using the Matlab function, which measures the distance of a data point from the mean in terms of the standard deviation. To estimate the z-scored amplitude, variance, and peak timing of the Down state onsets, we used a time window of 200 ms from 0-200 ms and fitted a Gaussian using Matlab. To determine the UP state onset amplitude, we calculated the maximum z-score values within a time window ranging from -200ms to -150ms before SWRs as well as from 150 ms to 200 ms after SWRs.

### Neural Decoding (Position Estimation) and Replay Detection

We conducted replay analysis using mice running in a linear maze measuring approximately 200 cm. The mice were subjected to a mild food-restricted protocol (5-6 grams of food/day), ensuring they did not lose weight during the running periods (5 days). They ran in the maze for approximately 30 min and were then placed back in their home cage, which was devoid of visual cues, and recorded their sleep for about 1-2 hours. To compute the joint probability distribution of position from neuronal firing activity, we employed a Bayesian reconstruction algorithm described by Zhang et al., 1998 and Davidson et al., 2009. This curve from CA1 units, as shown in Figure 5A1. For each unit, we constructed a joint tuning curve based on the linearized position (with 4 cm bins) using all the spikes emitted during the running session (Figure 5A1). To smooth the curve, we applied a Gaussian kernel with standard deviation of 6 cm. Interneurons with a mean firing rate greater than 9 Hz were excluded from the analysis. To estimate the position, we computed the marginal distribution of these estimates over position using 80% of the data as “training data.” We then validated the estimation using run epochs, as illustrated in Figure 5A2-3. During the testing period, we used the training models to estimate the position. To assess the reconstruction accuracy across the track, we calculated confusion matrices (Figure 5A2) and compared the maximum likelihood estimates of position with the mouse’s actual behavior to determine the median error (Figure 5A3) for the model.

Once we developed a model capable of estimating the position of mice with a median error accuracy of less than 20cm, our next step was to identify potential replay candidate events. We achieved this by using a smoothed histogram constructed from the multi-unit activity (MUA) of isolated unit clusters with 1 ms bins and a Gaussian kernel (SD = 25 ms) to smooth the signal. To detect candidate replay events, we applied a threshold of 3 standard deviations from the mean of the MUA. For an event to be classified as a replay candidate, it needed to last between 75 and 500 ms. We then binned the spikes from each candidate replay event at 5 ms intervals. Subsequently, we employed decoding methods to estimate the position within each candidate event.

To qualify as a “replay,” a candidate event had to exhibit temporal trajectories demonstrating both linear and nonlinear statistical dependencies, as described by Liu et al. (2018)(S. Liu, Grosmark, and Chen 2018). This involved fitting each event’s spatial trajectory using a linear model (Davidson, Kloosterman, and Wilson 2009; Wu and Foster 2014) as well as a weighted distance correlations model (S. Liu, Grosmark, and Chen 2018). To validate the significance of each candidate event, we conducted a statistical test. This involved comparing the correlation of the candidate event with the correlation of shuffled spiking activity in three distinct directions (row, column, and both) using Monte Carlo simulations. For assessing replays, we have provided the code at this link: (https://github.com/shizhaoliu/memory). Replays with a statistical p-value below 0.05 were included in the analysis.

### Statistical Analysis

All statistical analyses were done using robust ANOVA or t-test with 20% trimmed means and bootstrap-t. Here is a link to a GitHub repository containing the code and data used to perform the statistical tests: https://github.com/410pfeliciano/data_replay_coordination_to_rtc_pfc_states/tree/main (see Wilcox and Rousselet 2023; Mair and Wilcox 2020).

### Histology and Tetrode Localization

After completing the electrophysiological experiments, the mice were anesthetized using isoflurane until they were unresponsive to tail/toe pinches. Electrical currents were administered into each tetrode at 40 µA/tetrode for approximately 20 seconds. Subsequently, the mice were transcardially perfused with 1× PBS (30-40 mL) and then with 10% formalin at a rate of 1.3 mL/min until muscle contractions occurred. The brains were stored in 10% formalin for a minimum of 24 h before being sectioned using a vibratome. Coronal sections with a thickness of 50 µm were collected to ensure the accurate placement of all tetrodes.

## References

Ahmadi, Nur, Timothy G. Constandinou, and Christos-Savvas Bouganis. 2021. “Inferring Entire Spiking Activity from Local Field Potentials.” Scientific Reports 11 (1): 19045.

Aleman-Zapata, Adrian, Jacqueline van der Meij, and Lisa Genzel. 2022. “Disrupting Ripples: Methods, Results, and Caveats in Closed-Loop Approaches in Rodents.” Journal of Sleep Research 31 (6): e13532.

Aleman-Zapata, Adrian, Richard G. M. Morris, and Lisa Genzel. 2022. “Sleep Deprivation and Hippocampal Ripple Disruption after One-Session Learning Eliminate Memory Expression the next Day.” Proceedings of the National Academy of Sciences of the United States of America 119 (44): e2123424119.

Alexander, Andrew S., Lara M. Rangel, David Tingley, and Douglas A. Nitz. 2018. “Neurophysiological Signatures of Temporal Coordination between Retrosplenial Cortex and the Hippocampal Formation.” Behavioral Neuroscience 132 (5): 453–68.

Bakker, Hanna den, Marie Van Dijck, Jyh-Jang Sun, and Fabian Kloosterman. 2022. “Sharp-Wave Ripple Associated Activity in the Medial Prefrontal Cortex Supports Spatial Rule Switching.” bioRxiv. https://doi.org/10.1101/2022.11.03.515023.

Battaglia, Francesco P., Gary R. Sutherland, and Bruce L. McNaughton. 2004. “Hippocampal Sharp Wave Bursts Coincide with Neocortical ‘up-State’ Transitions.” Learning & Memory 11 (6): 697–704.

Buzsáki, G. 1989. “Two-Stage Model of Memory Trace Formation: A Role for ‘noisy’ Brain States.” Neuroscience. https://doi.org/10.1016/0306-4522(89)90423-5.

Buzsáki, György. 2015. “Hippocampal Sharp Wave-Ripple: A Cognitive Biomarker for Episodic Memory and Planning.” Hippocampus 25 (10): 1073–1188.

Carr, Margaret F., Shantanu P. Jadhav, and Loren M. Frank. 2011. “Hippocampal Replay in the Awake State: A Potential Substrate for Memory Consolidation and Retrieval.” Nature Neuroscience. https://doi.org/10.1038/nn.2732.

Chambers, Anna R., Christoffer Nerland Berge, and Koen Vervaeke. 2022. “Cell-Type-Specific Silence in Thalamocortical Circuits Precedes Hippocampal Sharp-Wave Ripples.” Cell Reports 40 (4): 111132.

Chen, Zhe, and Matthew A. Wilson. 2017. “Deciphering Neural Codes of Memory during Sleep.” Trends in Neurosciences 40 (5): 260–75.

Choi, Young-Seok, Matthew A. Koenig, Xiaofeng Jia, and Nitish V. Thakor. 2010. “Quantifying Time-Varying Multiunit Neural Activity Using Entropy Based Measures.” IEEE Transactions on Bio-Medical Engineering 57 (11). https://doi.org/10.1109/TBME.2010.2049266.

Choudhary, Krishna, Sven Berberich, Thomas T. G. Hahn, and Mayank R. Mehta. 2022. “Theory of Spontaneous Persistent Activity and Inactivity in Vivo Reveals Differential Cortico-Entorhinal Functional Connectivity.” bioRxiv. https://doi.org/10.1101/2022.04.15.488496.

Connelly, William M., Michael Laing, Adam C. Errington, and Vincenzo Crunelli. 2015. “The Thalamus as a Low Pass Filter: Filtering at the Cellular Level Does Not Equate with Filtering at the Network Level.” Frontiers in Neural Circuits 9: 89.

Davidson, Thomas J., Fabian Kloosterman, and Matthew A. Wilson. 2009. “Hippocampal Replay of Extended Experience.” Neuron 63 (4): 497–507.

Davoudi, Heydar, and David J. Foster. 2019. “Acute Silencing of Hippocampal CA3 Reveals a Dominant Role in Place Field Responses.” Nature Neuroscience 22 (3): 337–42.

De Filippo, Roberto, and Dietmar Schmitz. 2023. “Differential Ripple Propagation along the Hippocampal Longitudinal Axis.” eLife 12 (April). https://doi.org/10.7554/eLife.85488.

Ecker, András, Bence Bagi, Eszter Vértes, Orsolya Steinbach-Németh, Mária R. Karlócai, Orsolya I. Papp, István Miklós, et al. 2022. “Hippocampal Sharp Wave-Ripples and the Associated Sequence Replay Emerge from Structured Synaptic Interactions in a Network Model of Area CA3,” January. https://doi.org/10.7554/eLife.71850.

Egorov, Alexei V., Bassam N. Hamam, Erik Fransén, Michael E. Hasselmo, and Angel A. Alonso. 2002. “Graded Persistent Activity in Entorhinal Cortex Neurons.” Nature 420 (6912): 173–78.

Ego-Stengel, Valérie, and Matthew A. Wilson. 2010. “Disruption of Ripple-Associated Hippocampal Activity during Rest Impairs Spatial Learning in the Rat.” Hippocampus 20 (1): 1–10.

Eichenbaum, Howard. 2017. “Prefrontal–hippocampal Interactions in Episodic Memory.” Nature Reviews Neuroscience. https://doi.org/10.1038/nrn.2017.74.

Fanselow, Erika E., and Barry W. Connors. 2010. “The Roles of Somatostatin-Expressing (GIN) and Fast-Spiking Inhibitory Interneurons in UP-DOWN States of Mouse Neocortex.” Journal of Neurophysiology 104 (2): 596–606.

Fernández-Ruiz, Antonio, Azahara Oliva, Eliezyer Fermino de Oliveira, Florbela Rocha-Almeida, David Tingley, and György Buzsáki. 2019. “Long-Duration Hippocampal Sharp Wave Ripples Improve Memory.” Science 364 (6445): 1082–86.

Ferreira-Fernandes, Emanuel, Bárbara Pinto-Correia, Carolina Quintino, and Miguel Remondes. 2019. “A Gradient of Hippocampal Inputs to the Medial Mesocortex.” Cell Reports 29 (10): 3266–79.e3.

Foster, David J. 2017. “Replay Comes of Age.” Annual Review of Neuroscience 40 (July): 581–602.

Fries, Pascal. 2005. “A Mechanism for Cognitive Dynamics: Neuronal Communication through Neuronal Coherence.” Trends in Cognitive Sciences 9 (10): 474–80.

Girardeau, Gabrielle, Karim Benchenane, Sidney I. Wiener, György Buzsáki, and Michaël B. Zugaro. 2009. “Selective Suppression of Hippocampal Ripples Impairs Spatial Memory.” Nature Neuroscience 12 (10): 1222–23.

Groen, Thomas van, and J. Michael Wyss. 1990. “Connections of the Retrosplenial Granular a Cortex in the Rat.” The Journal of Comparative Neurology. https://doi.org/10.1002/cne.903000412.

Groen, T. van, and J. M. Wyss. 1992. “Connections of the Retrosplenial Dysgranular Cortex in the Rat.” The Journal of Comparative Neurology 315 (2): 200–216.

Hahn, Thomas T. G., James M. McFarland, Sven Berberich, Bert Sakmann, and Mayank R. Mehta. 2012. “Spontaneous Persistent Activity in Entorhinal Cortex Modulates Cortico-Hippocampal Interaction in Vivo.” Nature Neuroscience 15 (11): 1531–38.

Hale, Gregory (gregory John). 2015. “Timing and Hippocampal Information Processing.” Massachusetts Institute of Technology. http://hdl.handle.net/1721.1/100872.

Jackson, Jesse, Mahesh M. Karnani, Boris V. Zemelman, Denis Burdakov, and Albert K. Lee. 2018. “Inhibitory Control of Prefrontal Cortex by the Claustrum.” Neuron 99 (5): 1029–39.e4.

Jadhav, Shantanu P., Gideon Rothschild, Demetris K. Roumis, and Loren M. Frank. 2016. “Coordinated Excitation and Inhibition of Prefrontal Ensembles during Awake Hippocampal Sharp-Wave Ripple Events.” Neuron 90 (1): 113–27.

Ji, Daoyun, and Matthew A. Wilson. 2007. “Coordinated Memory Replay in the Visual Cortex and Hippocampus during Sleep.” Nature Neuroscience. https://doi.org/10.1038/nn1825.

Kaefer, Karola, Federico Stella, Bruce L. McNaughton, and Francesco P. Battaglia. 2022. “Replay, the Default Mode Network and the Cascaded Memory Systems Model.” Nature Reviews. Neuroscience 23 (10): 628–40.

Kajikawa, Koichiro, Brad K. Hulse, Athanassios G. Siapas, and Evgueniy V. Lubenov. 2022. “UP-DOWN States and Ripples Differentially Modulate Membrane Potential Dynamics across DG, CA3, and CA1 in Awake Mice.” eLife 11 (July). https://doi.org/10.7554/eLife.69596.

Karalis, Nikolaos, and Anton Sirota. 2022. “Breathing Coordinates Cortico-Hippocampal Dynamics in Mice during Offline States.” Nature Communications 13 (1): 467.

Karimi Abadchi, J., Mojtaba Nazari-Ahangarkolaee, Sandra Gattas, Edgar Bermudez-Contreras, Artur Luczak, Bruce L. McNaughton, and Majid H. Mohajerani. 2020. “Spatiotemporal Patterns of Neocortical Activity around Hippocampal Sharp-Wave Ripples.” eLife 9 (March). https://doi.org/10.7554/eLife.51972.

Kim, Jaekyung, Abhilasha Joshi, Loren Frank, and Karunesh Ganguly. 2023. “Cortical–hippocampal Coupling during Manifold Exploration in Motor Cortex.” Nature. https://doi.org/10.1038/s41586-022-05533-z.

Kitamura, Takashi, Sachie K. Ogawa, Dheeraj S. Roy, Teruhiro Okuyama, Mark D. Morrissey, Lillian M. Smith, Roger L. Redondo, and Susumu Tonegawa. 2017. “Engrams and Circuits Crucial for Systems Consolidation of a Memory.” Science 356 (6333): 73–78.

Klinzing, Jens G., Niels Niethard, and Jan Born. 2019. “Mechanisms of Systems Memory Consolidation during Sleep.” Nature Neuroscience. https://doi.org/10.1038/s41593-019-0467-3.

Kudrimoti, H. S., C. A. Barnes, and B. L. McNaughton. 1999. “Reactivation of Hippocampal Cell Assemblies: Effects of Behavioral State, Experience, and EEG Dynamics.” The Journal of Neuroscience: The Official Journal of the Society for Neuroscience 19 (10): 4090–4101.

Lee, Albert K., and Matthew A. Wilson. 2002. “Memory of Sequential Experience in the Hippocampus during Slow Wave Sleep.” Neuron 36 (6): 1183–94.

Liu, Shizhao, Andres D. Grosmark, and Zhe Chen. 2018. “Methods for Assessment of Memory Reactivation.” Neural Computation 30 (8): 2175–2209.

Liu, Xin, Chi Ren, Yichen Lu, Yixiu Liu, Jeong-Hoon Kim, Stefan Leutgeb, Takaki Komiyama, and Duygu Kuzum. 2021. “Multimodal Neural Recordings with Neuro-FITM Uncover Diverse Patterns of Cortical-Hippocampal Interactions.” Nature Neuroscience 24 (6): 886–96.

Logothetis, N. K., O. Eschenko, Y. Murayama, M. Augath, T. Steudel, H. C. Evrard, M. Besserve, and A. Oeltermann. 2012. “Hippocampal–cortical Interaction during Periods of Subcortical Silence.” Nature. https://doi.org/10.1038/nature11618.

MacPherson, Sarah E., and Sergio Della Sala. 2019. Cases of Amnesia: Contributions to Understanding Memory and the Brain. Routledge.

Maingret, Nicolas, Gabrielle Girardeau, Ralitsa Todorova, Marie Goutierre, and Michaël Zugaro. 2016. “Hippocampo-Cortical Coupling Mediates Memory Consolidation during Sleep.” Nature Neuroscience 19 (7): 959–64.

Mair, Patrick, and Rand Wilcox. 2020. “Robust Statistical Methods in R Using the WRS2 Package.” Behavior Research Methods 52 (2): 464–88.

McClelland, James L., Bruce L. McNaughton, and Randall C. O’Reilly. 1995. “Why There Are Complementary Learning Systems in the Hippocampus and Neocortex: Insights from the Successes and Failures of Connectionist Models of Learning and Memory.” Psychological Review 102 (3): 419–57.

Melzer, Sarah, and Hannah Monyer. 2020. “Diversity and Function of Corticopetal and Corticofugal GABAergic Projection Neurons.” Nature Reviews. Neuroscience 21 (9): 499–515.

Narikiyo, Kimiya, Rumiko Mizuguchi, Ayako Ajima, Momoko Shiozaki, Hiroki Hamanaka, Joshua P. Johansen, Kensaku Mori, and Yoshihiro Yoshihara. 2020. “The Claustrum Coordinates Cortical Slow-Wave Activity.” Nature Neuroscience 23 (6): 741–53.

Ngo, Hong-Viet, Juergen Fell, and Bernhard Staresina. 2020. “Sleep Spindles Mediate Hippocampal-Neocortical Coupling during Long-Duration Ripples.” eLife 9 (July). https://doi.org/10.7554/eLife.57011.

Nitzan, Noam, Sam McKenzie, Prateep Beed, Daniel Fine English, Silvia Oldani, John Jan Tukker, György Buzsáki, and Dietmar Schmitz. 2020. “Propagation of Hippocampal Ripples to the Neocortex by Way of a Subiculum-Retrosplenial Pathway.” Nature Communications 11 (1947). https://doi.org/10.1038/s41467-020-15787-8.

Nitzan, Noam, Rachel Swanson, Dietmar Schmitz, and György Buzsáki. 2022. “Brain-Wide Interactions during Hippocampal Sharp Wave Ripples.” Proceedings of the National Academy of Sciences of the United States of America 119 (20): e2200931119.

Oliva, Azahara, Antonio Fernández-Ruiz, György Buzsáki, and Antal Berényi. 2016. “Role of Hippocampal CA2 Region in Triggering Sharp-Wave Ripples.” Neuron 91 (6): 1342–55.

Opalka, Ashley N., Wen-Qiang Huang, Jun Liu, Hualou Liang, and Dong V. Wang. 2020. “Hippocampal Ripple Coordinates Retrosplenial Inhibitory Neurons during Slow-Wave Sleep.” Cell Reports 30 (2): 432–41.e3.

Opalka, Ashley N., and Dong V. Wang. 2020. “Hippocampal Efferents to Retrosplenial Cortex and Lateral Septum Are Required for Memory Acquisition.” Learning & Memory 27 (8): 310–18.

Pavlides, C., and J. Winson. 1989. “Influences of Hippocampal Place Cell Firing in the Awake State on the Activity of These Cells during Subsequent Sleep Episodes.” The Journal of Neuroscience: The Official Journal of the Society for Neuroscience 9 (8): 2907–18.

Pedrosa, Rafael, Mojtaba Nazari, Loig Kergoat, Christophe Bernard, Majid Mohajerani, Federico Stella, and Francesco Battaglia. n.d. “Hippocampal Ripples Coincide with ‘up-State’ and Cortical Spindles in Retrosplenial Cortex.” https://doi.org/10.1101/2022.12.19.521088.

Pedrosa, Rafael, Mojtaba Nazari, Majid H. Mohajerani, Thomas Knöpfel, Federico Stella, and Francesco P. Battaglia. 2022. “Hippocampal Gamma and Sharp Wave/ripples Mediate Bidirectional Interactions with Cortical Networks during Sleep.” Proceedings of the National Academy of Sciences of the United States of America 119 (44): e2204959119.

Penagos, Hector, Carmen Varela, and Matthew A. Wilson. 2017. “Oscillations, Neural Computations and Learning during Wake and Sleep.” Current Opinion in Neurobiology 44 (June): 193–201.

Peyrache, Adrien, Mehdi Khamassi, Karim Benchenane, Sidney I. Wiener, and Francesco P. Battaglia. 2009. “Replay of Rule-Learning Related Neural Patterns in the Prefrontal Cortex during Sleep.” Nature Neuroscience 12 (7): 919–26.

Prida, Liset M. de la. 2020. “Potential Factors Influencing Replay across CA1 during Sharp-Wave Ripples.” Philosophical Transactions of the Royal Society of London. Series B, Biological Sciences 375 (1799): 20190236.

Qin, Y. L., B. L. McNaughton, W. E. Skaggs, and C. A. Barnes. 1997. “Memory Reprocessing in Corticocortical and Hippocampocortical Neuronal Ensembles.” Philosophical Transactions of the Royal Society of London. Series B, Biological Sciences 352 (1360): 1525–33.

Reimann, Michael W., Costas A. Anastassiou, Rodrigo Perin, Sean L. Hill, Henry Markram, and Christof Koch. 2013. “A Biophysically Detailed Model of Neocortical Local Field Potentials Predicts the Critical Role of Active Membrane Currents.” Neuron 79 (2): 375–90.

Rothschild, Gideon, Elad Eban, and Loren M. Frank. 2017. “A Cortical-Hippocampal-Cortical Loop of Information Processing during Memory Consolidation.” Nature Neuroscience 20 (2): 251–59.

Saleem, Aman B., Paul Chadderton, John Apergis-Schoute, Kenneth D. Harris, and Simon R. Schultz. 2010. “Methods for Predicting Cortical UP and DOWN States from the Phase of Deep Layer Local Field Potentials.” Journal of Computational Neuroscience 29 (1-2): 49–62.

Sawangjit, Anuck, Carlos N. Oyanedel, Niels Niethard, Carolina Salazar, Jan Born, and Marion Inostroza. 2018. “The Hippocampus Is Crucial for Forming Non-Hippocampal Long-Term Memory during Sleep.” Nature 564 (7734): 109–13.

Siapas, Athanassios G., and Matthew A. Wilson. 1998. “Coordinated Interactions between Hippocampal Ripples and Cortical Spindles during Slow-Wave Sleep.” Neuron. https://doi.org/10.1016/s0896-6273(00)80629-7.

Sirota, Anton, Jozsef Csicsvari, Derek Buhl, and György Buzsáki. 2003. “Communication between Neocortex and Hippocampus during Sleep in Rodents.” Proceedings of the National Academy of Sciences of the United States of America 100 (4): 2065–69.

Skaggs, W. E., and B. L. McNaughton. 1996. “Replay of Neuronal Firing Sequences in Rat Hippocampus during Sleep Following Spatial Experience.” Science 271 (5257): 1870–73.

Sosa, Marielena, Hannah R. Joo, and Loren M. Frank. 2020. “Dorsal and Ventral Hippocampal Sharp-Wave Ripples Activate Distinct Nucleus Accumbens Networks.” Neuron 105 (4): 725–41.e8.

Stickgold, Robert. 2005. “Sleep-Dependent Memory Consolidation.” Nature 437 (7063): 1272–78.

Tang, Wenbo, Justin D. Shin, Loren M. Frank, and Shantanu P. Jadhav. 2017. “Hippocampal-Prefrontal Reactivation during Learning Is Stronger in Awake Compared with Sleep States.” The Journal of Neuroscience: The Official Journal of the Society for Neuroscience 37 (49): 11789–805.

Tingley, David, and Adrien Peyrache. 2020. “On the Methods for Reactivation and Replay Analysis.” Philosophical Transactions of the Royal Society of London. Series B, Biological Sciences 375 (1799): 20190231.

Todorova, Ralitsa, and Michaël Zugaro. 2019. “Isolated Cortical Computations during Delta Waves Support Memory Consolidation.” Science 366 (6463): 377–81.

Tomé Douglas Feitosa, Sadra Sadeh, and Claudia Clopath. 2022. “Coordinated Hippocampal-Thalamic-Cortical Communication Crucial for Engram Dynamics underneath Systems Consolidation.” Nature Communications 13 (1): 840.

Tse, Dorothy, Rosamund F. Langston, Masaki Kakeyama, Ingrid Bethus, Patrick A. Spooner, Emma R. Wood, Menno P. Witter, and Richard G. M. Morris. 2007. “Schemas and Memory Consolidation.” Science 316 (5821): 76–82.

Vaasjo, Lee O., Xiao Han, Abbigail N. Thurmon, Alina S. Tiemroth, Hallie Berndt, Madelyn Korn, Alexandra Figueroa, Rosa Reyes, Pedro A. Feliciano-Ramos, and Maria J. Galazo. 2022. “Characterization and Manipulation of Corticothalamic Neurons in Associative Cortices Using Syt6-Cre Transgenic Mice.” The Journal of Comparative Neurology 530 (7): 1020–48.

Varela, Carmen, and Matthew A. Wilson. 2020. “mPFC Spindle Cycles Organize Sparse Thalamic Activation and Recently Active CA1 Cells during Non-REM Sleep.” eLife 9 (June). https://doi.org/10.7554/eLife.48881.

Vyazovskiy, Vladyslav V., Umberto Olcese, Erin C. Hanlon, Yuval Nir, Chiara Cirelli, and Giulio Tononi. 2011. “Local Sleep in Awake Rats.” Nature. https://doi.org/10.1038/nature10009.

Wang, Dong V., and Satoshi Ikemoto. 2016. “Coordinated Interaction between Hippocampal Sharp-Wave Ripples and Anterior Cingulate Unit Activity.” The Journal of Neuroscience: The Official Journal of the Society for Neuroscience 36 (41): 10663–72.

Wierzynski, Casimir M., Evgueniy V. Lubenov, Ming Gu, and Athanassios G. Siapas. 2009. “State-Dependent Spike-Timing Relationships between Hippocampal and Prefrontal Circuits during Sleep.” Neuron 61 (4): 587–96.

Wilcox, Rand R., and Guillaume A. Rousselet. 2023. “An Updated Guide to Robust Statistical Methods in Neuroscience.” Current Protocols 3 (3): e719.

Wilson, Matthew A., and Bruce L. McNaughton. 1994. “Reactivation of Hippocampal Ensemble Memories During Sleep.” Science. https://doi.org/10.1126/science.8036517.

Wu, Xiaojing, and David J. Foster. 2014. “Hippocampal Replay Captures the Unique Topological Structure of a Novel Environment.” The Journal of Neuroscience: The Official Journal of the Society for Neuroscience 34 (19): 6459–69.

Wyass, J. Michael, J. Michael Wyass, and Thomas Van Groen. 1992. “Connections between the Retrosplenial Cortex and the Hippocampal Formation in the Rat: A Review.” Hippocampus. https://doi.org/10.1002/hipo.450020102.

Yamamoto, Jun, and Susumu Tonegawa. 2017. “Direct Medial Entorhinal Cortex Input to Hippocampal CA1 Is Crucial for Extended Quiet Awake Replay.” Neuron 96 (1): 217–27.e4.

Yamawaki, Naoki, Xiaojian Li, Laurie Lambot, Lynn Y. Ren, Jelena Radulovic, and Gordon M. G. Shepherd. 2019. “Long-Range Inhibitory Intersection of a Retrosplenial Thalamocortical Circuit by Apical Tuft-Targeting CA1 Neurons.” Nature Neuroscience 22 (4): 618–26.

Yoshida, Motoharu, Erik Fransén, and Michael E. Hasselmo. 2008. “mGluR-Dependent Persistent Firing in Entorhinal Cortex Layer III Neurons.” The European Journal of Neuroscience 28 (6): 1116–26.

Zucca, Stefano, Giulia D’Urso, Valentina Pasquale, Dania Vecchia, Giuseppe Pica, Serena Bovetti, Claudio Moretti, et al. 2017. “An Inhibitory Gate for State Transition in Cortex.” eLife 6 (May). https://doi.org/10.7554/eLife.26177.

