## Supplementary Figures for "Hippocampal memory reactivation during sleep is correlated with specific cortical states of the Retrosplenial and Prefrontal Cortices"

### Supplementary Figures
### Supplementary Figure 1A

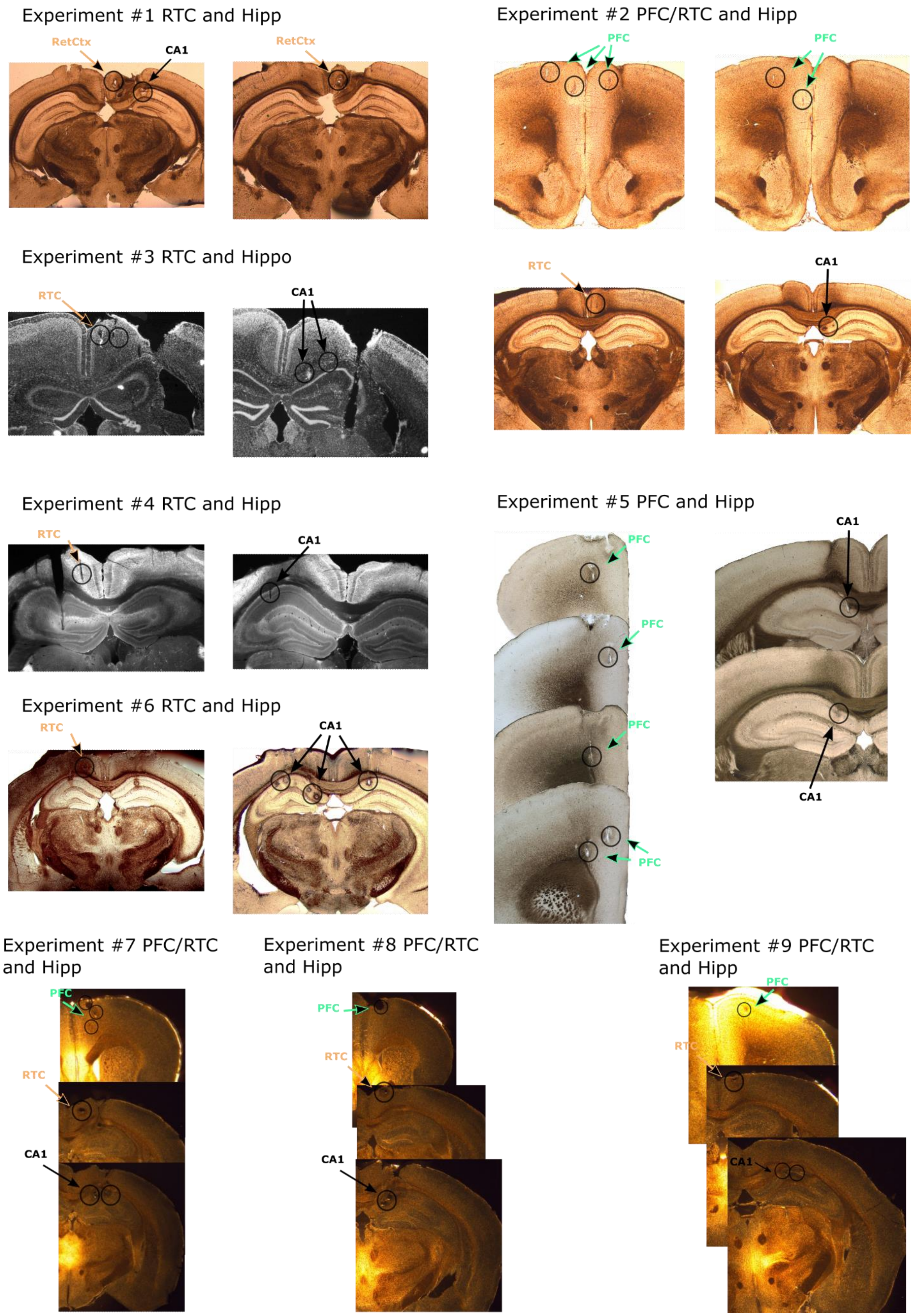


**Supplementary Figure 1A: post hoc Histology for the localization of chronically implanted tetrodes.** The circles represent the estimated location of the tip of a tetrode post hoc histology.

### Supplementary Figure 1B

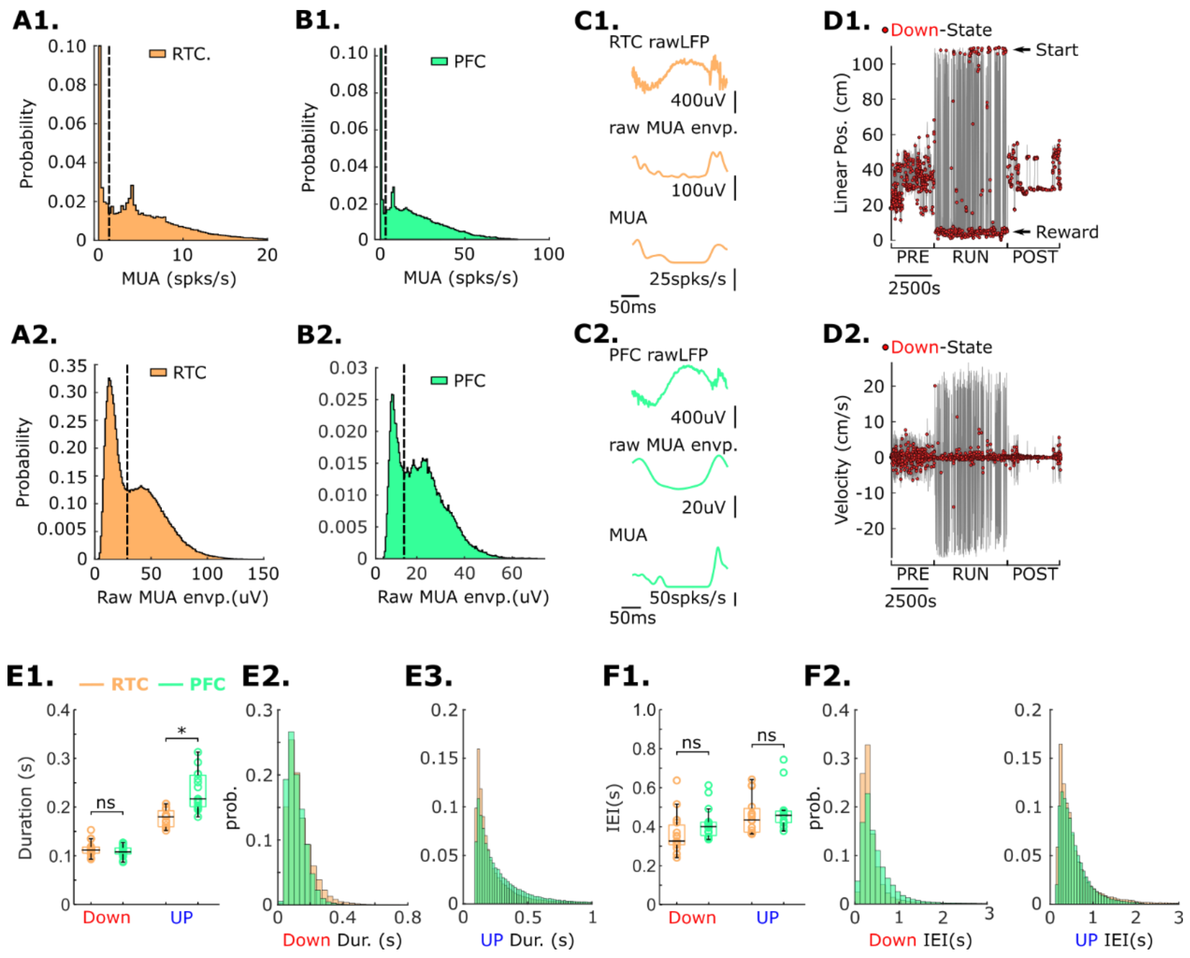


**Supplementary Figure 1B: Characteristics of Down and UP Cortical states in the RTC and PFC. A-B,** Bimodal distributions of multiunit activity (MUA) (A1-B1) and raw MUA envelop (A2-B2) for the RTC and PFC during sleep. We use the pit of the bimodal distribution as a threshold (black line) approximation for detecting cortical Down and UP states. **C1-2,** Single examples of detected cortical Down states in the RTC and PFC. **D1-2,** Cortical Down states occur during immobility. D1, The linearized position of a single experiment. Red circles represent a detected Down-state. D2, The velocity corresponding to D1. **E,** RTC, and PFC median UP and Down states duration distribution and cumulative probability. E1, RTC median Down- and UP-state durations were 0.112 (Down, IQR 0.106-0.117) and 0.180 (UP, IQR 0.161-0.189). For PFC was 0.108 (Down, IQR 0.104-0.116) and 0.217 (UP, IQR 0.202-0.263). Down-state duration RTC vs PFC p=0.601 95%CI[-0.011 0.014] and for UP-states duration RTC vs PFC p=0.04 95%CI[-0.051 -0.104]; robust ANOVA, Multiple comparisons 20% trimmed mean with bootstrap-t. **F,** RTC, and PFC median Down states inter-Event-interval(IEI) distribution and cumulative probability. F1, RTC median Down- and UP-state IEI were 0.326 (Down, IQR 0.308-0.396) and 0.434 (UP, IQR 0.375-0.486). For PFC was 0.401 (Down, IQR 0.360-0.422) and 0.458 (IQR 0.422-0.479). Down-state IEI RTC vs PFC p=0.155 95%CI[-0.129 0.019] and for UP-states IEI RTC vs PFC p=0.062 95%CI[-0.094 0.062]; robust ANOVA, post hoc 20% trimmed mean with bootstrap-t.

### Supplementary Figure 3

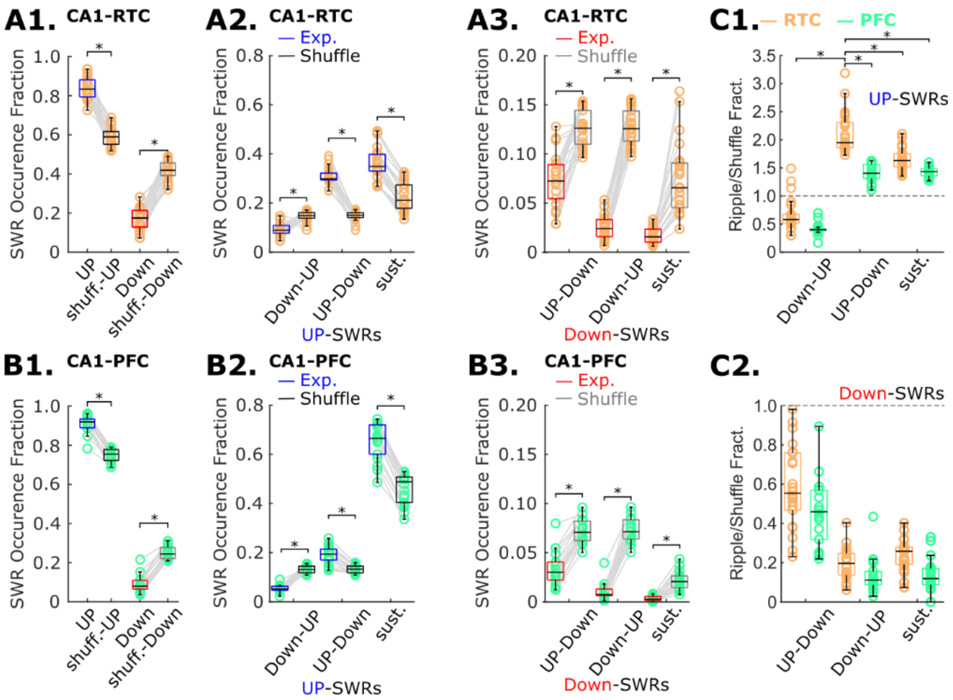


**Supplementary Figure 3: Comparison of cortico-hippocampal activity among hippocampal CA1, RTC, and PFC with their controls (SWRs timing shuffle). A1-B1,** Fraction of SWRs that occurred during a cortical UP- or Down-state of the RTC(A1) or PFC(B1). A1, RTC UP vs UP-shuffle, *p*=0 95%CI [0.202 0.302]; RTC Down vs Down-shuffle, *p*=0 95%CI [-0.302 -0.202]. B1, PFC UP vs UP-shuffle *p*=0 95%CI [0.134 0.194], PFC Down vs Down-shuffle *p*=0 95%CI [-0.194 -0.134]. **A2-B2,** Fraction of SWRs that occurred during configurations of the UP-state of the RTC(A2) or PFC(B2). RTC UP sust. exp. = 0.349(IQR: 0.331-0.398) vs sust. shuff. = 0.212(IQR: 0.179-0.271) (*p*=0 95%CI [0.089 0.191]); PFC UP sust. exp. = 0.667(IQR: 0.615-0.721) vs UP sust. shuff. = 0.49(IQR 0.411-0.509) (*p*=0 95%CI [0.121 0.274]). UP-Down, RTC exp. 0.299(IQR: 0.294-0.321) vs shuff. 0.151(IQR: 0.142-0.161)(p=0 95%CI [0.134 0.173]); PFC exp. 0.195(IQR: 0.170-.215) vs shuff. 0.133(IQR: 0.121-0.145)(*p*=0 95%CI [0.029 0.088]). Down-UP, RTC exp. 0.091(IQR: 0.079-0.109) vs shuff. 0.149(IQR: 0.139-0.163)(p=0 95%CI [-0.076 -0.041]) PFC exp. 0.052(IQR: 0.047-0.063) vs shuff. 0.132(IQR: 0.121-0.145)(p=0 95%CI [-0.076 -0.041]). **A3-B3,** Fraction of SWRs that occurred during configurations of the Down-state of the RTC(A2) or PFC(B2). RTC Down sust. exp. = 0.016(IQR: 0.010-0.023) vs sust. shuff. = 0.066(IQR: 0.046-0.090) (*p*=0 95%CI [0.121 0.274]); PFC Down sust. exp. = 0.003(IQR: 0.001-0.005) vs Down sust. shuff. = 0.021(IQR: 0.015-0.026) (*p*=0 95%CI [-0.072 -0.032]). Down-UP, RTC exp. 0.024(IQR: 0.016-0.033) vs shuff. 0.126(IQR: 0.113-0.142)(*p*=0 95%CI [0.029 0.088]); PFC exp. 0.024(IQR: 0.016-0.033) vs shuff. 0.126(IQR: 0.113-0.142)(*p*=0 95%CI [-0.119 -0.086]). UP-Down, RTC exp. 0.052(IQR: 0.046-0.063) vs shuff. 0.132(IQR: 0.121-0.145)(*p*=0 95%CI [-0.094 -0.063]) PFC exp. 0.072(IQR: 0.056-0.088) vs shuff. 0.126(IQR: 0.110-0.141)(*p*=0 95%CI [-0.073 -0.032]). **C1-C2**, Ratio of exp./shuff for UP(C1) and Down(C2) configurations. C2, RTC UP-Down vs RTC Down-UP, PFC Down-UP, PFC UP-Down, RTC sust. UP, PFC sust. UP, *p*=0 95%CI [-1.672 -1.198], *p*=0 95%CI [-1.845 -1.427], *p*=0 95%CI [-0.839 -0.375], *p*=0 95%CI [-3.891 -3.436], *p*=0 95%CI [-3.679 -3.249], respectively.

### Supplementary Figure 3B

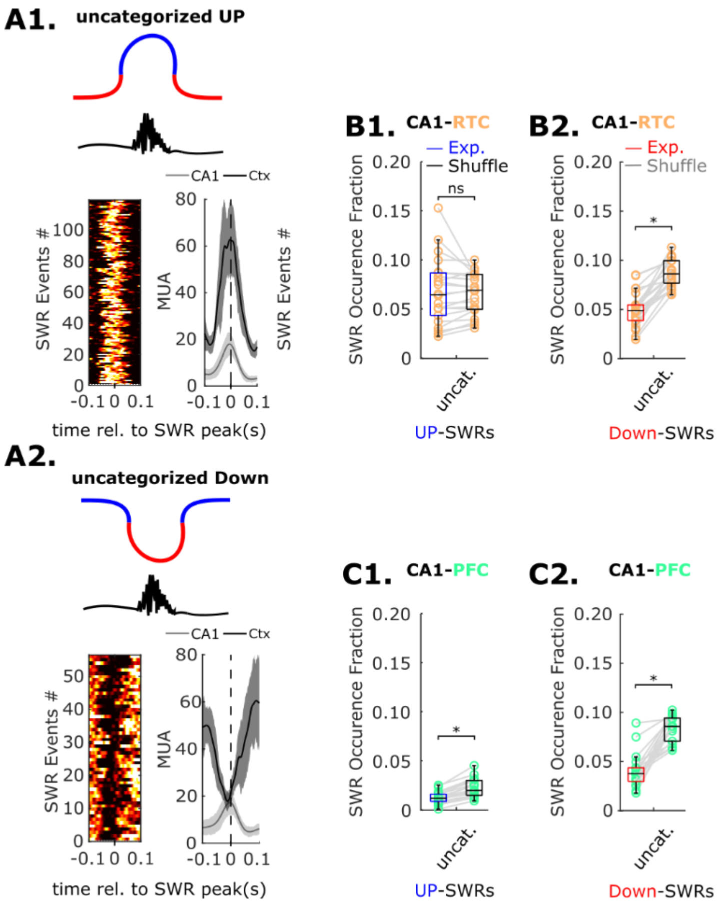


**Supplementary Figure 3B: Uncategorized SWRs during UP and Down-states.** These were SWRs that occurred between Down-states but during a UP state(A1, B1, C1) or between UP states during a Down state(A2, B2, C2). Given the brief duration of the UP- or Down-state events in these configurations, it was challenging to determine if they were associated with Delta or SO. Consequently, we decided to exclude them from our analysis. **A1,** Cartoon illustration and multiunit activity from cortex and hippocampus for SWRs classified as uncategorized UP. **B1 and C1,** Fraction of Uncategorized UP SWRs for RTC(B1) and PFC(C1) with their respective shuffles. **A2,** Cartoon illustration and multiunit activity from cortex and hippocampus for SWRs classified as uncategorized Down. **B2 and C2,** Fraction of Uncategorized Down SWRs for RTC(B2) and PFC(C2) with their respective shuffles.

### Supplementary Figure 4

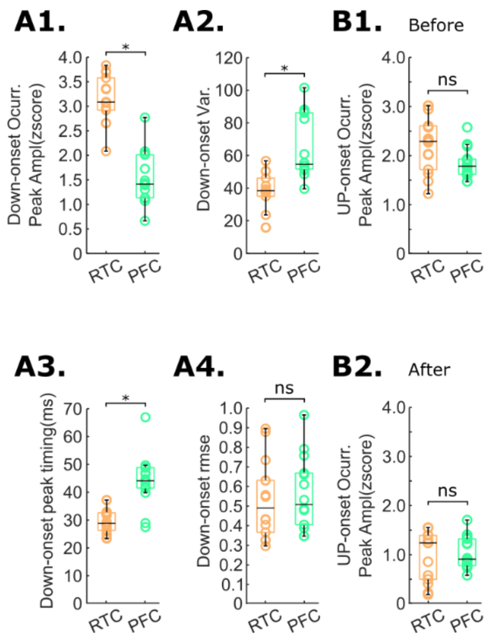


**Supplementary Figure 4: Peri-event histograms analysis for the onset of cortical Down and UP states of the RTC and PFC. A,** Gaussian fit analysis for the onset of Down states. Gaussian fit estimation of the peak amplitude (A1), variance (A2), timing (A3), and root-mean-square deviation (A4) for the onset of Down states. **B,** Peak amplitude before (B1) and after (B2) for onset of cortical UP states of the RTC and PFC. *= $p$<0.01, ns= $p$>0.05
